# Circulation of West Nile Virus and Usutu Virus in Birds in Germany, 2021 and 2022

**DOI:** 10.1101/2024.08.28.610071

**Authors:** Franziska Schopf, Balal Sadeghi, Felicitas Bergmann, Dominik Fischer, Ronja Rahner, Kerstin Müller, Anne Günther, Anja Globig, Markus Keller, Rebekka Schwehn, Vanessa Guddorf, Maximilian Reuschel, Luisa Fischer, Oliver Krone, Monika Rinder, Karolin Schütte, Volker Schmidt, Kristin Heenemann, Anne Schwarzer, Christine Fast, Carola Sauter-Louis, Christoph Staubach, Renke Lühken, Jonas Schmidt-Chanasit, Florian Brandes, Michael Lierz, Rüdiger Korbel, Thomas W. Vahlenkamp, Martin H. Groschup, Ute Ziegler

**Affiliations:** Friedrich-Loeffler-Institut, Federal Research Institute for Animal Health, Institute of Novel and Emerging Infectious Diseases, 17493 Greifswald-Insel Riems, Germany; (F.S.); (B.S.); (F.B.); (M.K.); (A.S.); (C.F.); (M.H.G.); German Center of Infection Research (DZIF), Partner site Hamburg-Lübeck-Borstel-Riems, 17493 Greifswald-Insel Riems, Germany; Clinic for Birds, Reptiles, Amphibians and Fish, Justus Liebig University Giessen, 35392 Giessen, Germany; (D.F.); (R.R.); (M.L.); Der Grüne Zoo Wuppertal, 42117 Wuppertal, Germany; Department of Veterinary Medicine, Small Animal Clinic, Freie Universität Berlin, 14163 Berlin, Germany; (K.M.); Friedrich-Loeffler-Institut, Federal Research Institute for Animal Health, Institute of Diagnostic Virology, 17493 Greifswald-Insel Riems, Germany; (A.G.); Friedrich-Loeffler-Institut, Federal Research Institute for Animal Health, Institute of International Animal Health/One Health, 17493 Greifswald-Insel Riems, Germany; (A.Gl.); Department of Small Mammal, Reptile and Avian Diseases, University of Veterinary Medicine Hannover, Foundation, 30559 Hannover, Germany; (M.R.); (R.S.); Seehundstation Nationalpark-Haus Norden-Norddeich, 26506 Norden, Germany; (V.G.); Wildlife Research Institute, State Agency for Nature, Environment and Consumer Protection North Rhine-Westphalia, 53229 Bonn, Germany; (L.F.); Leibniz Institute for Zoo and Wildlife Research (IZW), Department of Wildlife Diseases, 10315 Berlin, Germany; (O.K.); Clinic for Birds, Small Mammals, Reptiles and Ornamental Fish, Centre for Clinical Veterinary Medicine, Ludwig Maximilians University Munich, 85764 Oberschleißheim, Germany; (M.Ri.); (R.K.); Wildlife Rescue and Conservation Centre, 31553 Sachsenhagen, Germany; (K.S.); (F.Br.); Clinic for Birds and Reptiles, Faculty of Veterinary Medicine, Leipzig University, 04103 Leipzig, Germany; (V.S.); Institute of Virology, Faculty of Veterinary Medicine, Leipzig University, 04103 Leipzig, Germany; (K.H.); (T.W.V.); Friedrich-Loeffler-Institut, Federal Research Institute for Animal Health, Institute of Epidemiology, 17493 Greifswald-Insel Riems, Germany; (C.S.-L.); (C.S.); Bernhard Nocht Institute for Tropical Medicine, Department of Arbovirology and Entomology, 20359 Hamburg, Germany; (R.L.); (J.S.-C.)

**Keywords:** arbovirus/flavivirus, zoonosis, monitoring network, phylogeny, serology, Europe

## Abstract

**Background:** Usutu virus (USUV) and West Nile virus (WNV) are zoonotic arthropod-borne orthoflaviviruses. The enzootic transmission cycles of both include *Culex* mosquitoes as vectors and birds as amplifying hosts. For more than ten years, these viruses have been monitored in birds in Germany by a multidisciplinary network. While USUV is present nationwide, WNV used to be restricted to the central-east.

**Methods:** In 2021 and 2022, over 2300 live bird blood samples and organs from over 3000 deceased birds were subjected to molecular and serological analysis regarding presence of WNV and USUV. The samples were collected at sites all over Germany.

**Results:** Circulation of both viruses increased in 2022. For USUV, the nationwide presence of lineages Africa 3 and Europe 3 reported in previous years was confirmed. Lineage Europe 2, formerly restricted to the German east, was able to expand westward. Nonetheless, USUV neutralizing antibody (nAb) detection rates remained low (< 9%). 2021 and 2022 were characterized by stable enzootic circulation of WNV lineage 2, dominated by one previously identified subcluster (95% of generated sequences). In 2022, more than 20% of birds in the endemic region in eastern Germany carried nAb against WNV. Serological data also indicate expanding WNV circulation west and south of the known hotspots in Germany.

**Conclusions:** USUV circulates enzootically nationwide. Emergence of WNV at several new locations in Germany with a potential increase in human infections may be imminent. In this context, wild bird monitoring serves as a capable early-warning system in a One Health setting.

## 1 Introduction

West Nile virus (WNV, *Orthoflavivirus nilense (International Committee on Taxonomy of Viruses 2024)*) is an important zoonotic arthropod-borne virus and almost ubiquitously prevalent in the world. It is transmitted between mosquitoes and local birds (Hayes et al. 2005; Brault 2009). In this enzootic cycle *Culex* sp. are the most important vectors and reservoirs, while notably bird species from the orders Passeriformes, Accipitriformes, Falconiformes and Strigiformes are relevant amplifiers for WNV (Brault 2009; Ciota 2017). The course of the infection wave depends on the susceptibility of the local bird population, but also on vector competence and availability associated with environmental conditions (Kilpatrick et al. 2007; Kilpatrick 2011). If the combination of these factors is favourable to WNV transmission, it becomes an important zoonosis, where waves of infection in humans as dead-end hosts typically follow the cases of illness in birds (Petersen et al. 2003). As horses also serve as dead-end hosts, they can likewise develop clinical illness with neurological symptoms potentially resulting in encephalitis with fatal outcome (Gardner et al. 2007).

Over the last years infections with WNV in birds, horses and humans have increasingly become the focus of Central Europe due to climate change (Camp and Nowotny 2020). While WNV infections occur only sporadically in some neighbouring countries of Germany so far (e.g. the Netherlands or Poland), the virus has been circulating in Germany since 2018 with annually recurring noticeable cases of disease in birds, horses and humans (Ziegler et al. 2019b). This was likely favoured by extraordinarily high environmental temperatures in 2018 and 2019, which allowed a low extrinsic incubation period in competent mosquitoes resulting in a quicker virus transmission cycle and drove the WNV epizootic emergence in Germany during the succeeding years (Ziegler et al. 2020).

It is interesting to note that WNV genomes have nearly exclusively been detected in the central and eastern part of Germany so far (Ziegler et al. 2019b; Ziegler et al. 2020; Santos et al. 2023). This correlates with a high WNV seroprevalence in resident birds of 14.77% (2019) and 16.15% (2020), in addition to a noticeable tendency for a westward and southward expansion (Ziegler et al. 2022).

The phylogenetic analyses for WNV revealed that the causative strain in Germany belonged to the central European subclade II (Ziegler et al. 2019b). It could be demonstrated that Germany experienced several WNV introduction events and that strains of the Eastern German clade, which was introduced in a single event, dominated the virus variants in 2018 and 2019 (Ziegler et al. 2020). A new WNV phylogenetic study for the year 2020 revealed that the genetic diversity of the WNV population in Germany is characterized by the sole enzootic maintenance of one dominant WNV subcluster (Santos et al. 2023).

While migratory birds play an important role in the introduction of viruses into new areas (Reiter 2010), the detection of pathogens in local birds indicates the presence of these infectious agents in a defined area. In this context, the German wild bird-associated zoonoses network (WBA-Zoo) stands as a nationwide long-lived and well-functioning surveillance structure for arboviruses with various avifaunal experts as cooperation partners. It forms the basis for such monitoring and has proven its capabilities in the past (Seidowski et al. 2010; Ziegler et al. 2012; Ziegler et al. 2015; Michel et al. 2018; Michel et al. 2019; Ziegler et al. 2022).

The WBA-Zoo has contributed to the timely recognition of WNV entry into Germany and to the regional mapping of its annual spread. The early detection of Usutu virus (USUV; *Orthoflavivirus usutuense (International Committee on Taxonomy of Viruses 2024)*) entry into the bird population in Germany more than 10 years ago stands as an important proof of principle for this network (Bergmann et al. 2023a).

USUV is closely related with and similar to WNV as it circulates in approximately the same enzootic cycle between susceptible birds and competent mosquitoes and can also be transmitted to mammals by potential bridge-vectors. It possesses lower zoonotic potential than WNV, but nonetheless more than 100 cases of human infection with USUV were reported in the European Union / European Economic Area between 2012 and 2021, 12 of which were associated with neurological symptoms (Nikolay 2015; Zannoli and Sambri 2019; Angeloni et al. 2023). USUV has been associated with mass mortalities of various wild and zoo bird species, preferentially Eurasian blackbirds (*Turdus merula*) or great grey owls (*Strix nebulosa*), since its introduction into Europe in 1996 or earlier (Weissenböck et al. 2013). The virus rapidly expanded throughout Europe, reaching Germany in 2010/2011 with the first detections in *Culex* sp. mosquitoes and the bird population, respectively (Jöst et al. 2011; Becker et al. 2012). Initial phylogenetic studies revealed the presence and progressive evolution of different USUV lineages in Germany, with emphasis on USUV lineages Europe 3 and Africa 3 (Bergmann et al. 2023a). Furthermore, USUV lineages Europe 2 and Africa 2 were detected in central-eastern parts of the country. The USUV seroprevalence in resident birds in the different regions varied depending on the infection events and years (Michel et al. 2018; Michel et al. 2019; Ziegler et al. 2022).

The monitoring study presented here builds on the results of previous years. It illustrates the molecular and serological data of the German WBA-Zoo for the years 2021 and 2022, highlighting similarities and differences in USUV and WNV circulation e.g. regarding geographical distribution and affected bird species. The phylogenetic developments in the occurrence of these two viruses in Germany within this time span will also be examined in more detail.

## 2 Materials and Methods

### 2.1 Sample Collection

The WBA-Zoo boasts a very respectable number of cooperation partners from the avifaunal community. As previously described (Michel et al. 2019), this results in sample collection sites spread widely across Germany.

#### 2.1.1 Blood samples (live bird monitoring)

The first monitoring panel of this study consists of blood samples taken either from diseased birds in order to perform routine haematological as well as chemical analyses or from freshly euthanized animals. The blood was drawn from the *Vena* (V.) *ulnaris*, *V. metatarsalis dorsalis* or the *V. jugularis externa dextra* and in euthanized birds often intracardially. The cooperation partners centrifuged the blood sample on site and separated serum from the coagulated blood portion (cruor). Bird identification methods such as documentation of ring numbers, weighing as well as morphologic and radiographic examination helped to minimize the risk of double sampling. The samples were picked up from the sampling sites in regular intervals and catalogued at the national reference laboratory (NRL) for WNV at the Federal Research Institute for Animal Health (Friedrich-Loeffler-Institut, FLI) in Germany. Cruor and serum were stored until further analysis at −70°C and −20°C, respectively.

More than 2350 blood samples were taken and processed for this study, 1299 from the year 2021 and 1057 from 2022. The sampled birds belong to 19 different taxonomic orders (Table 1). Every species was categorized according to its (non-) migratory behaviour. Resident wild birds as well as captive birds in zoos or sentinel holdings remain in their habitat throughout the year. Migratory birds can be differentiated into partial migrants (some members of the population leaving their German habitat, others staying behind), short-distance migrants (traveling up to 2000 km but not crossing the Sahara Desert) and long-distance migrants (covering distances of 3000-4000 km and/or crossing the Sahara Desert). All avian species, their migration pattern and the corresponding number of tested birds from each region can be viewed in detail in Supplementary Tables S1-S3.

**Table 1:** Numbers of wild bird blood samples from the live bird panel itemized according to year of sampling and taxonomic order of the species of origin.

| **Taxonomic order** | **2021** | **2022** | **Total** |
| --- | --- | --- | --- |
| **Passeriformes** | 369 | 233 | 602 |
| **Accipitriformes** | 265 | 306 | 571 |
| **Falconiformes** | 79 | 53 | 132 |
| **Strigiformes** | 46 | 30 | 76 |
| **Anseriformes** | 75 | 42 | 117 |
| **Columbiformes** | 284 | 227 | 511 |
| **Charadriiformes** | 87 | 70 | 157 |
| **Ciconii-**<br>**/Pelecaniformes** | 21 | 13 | 34 |
| **Piciformes** | 52 | 32 | 84 |
| **Suliformes** | 1 | 34 | 35 |
| **Other** | 20 | 17 | 37 |
| **Total** | 1299 | 1057 | 2356 |

For the majority of blood samples, the cooperation partners additionally identified the corresponding wild bird as being adult or juvenile. These metadata can be used to divide the sample panel into three age groups. Firstly, birds that have most likely experienced less than two complete WNV/USUV transmission seasons. Secondly, birds that have most likely experienced more than one such season. Finally, birds for whom the age is unknown. In this study these three groups are labelled “juvenile”, “adult” and “age unknown”, respectively. There is a considerable temporal overlap between the first two subsets due to the high species diversity in our sample panel. This diversity results in a potentially great variation in length of life between birds to whom the same developmental denominator is applied. Some juvenile birds, mainly pigeon sp. (*Columba sp.*), were at times additionally labelled “pullus” by our cooperation partners. This term describes a non-fledged bird in its first set of feathers (dunes). In pigeons, this terminus was applied to birds until approximately 35 days post-hatch.

To guarantee continuity and comparability with results from previous years, we adhered to the previously published subdivision of Germany into three geographical regions (Ziegler et al. 2022). The northern and central-western parts of the country are labelled region A, the eastern and central eastern ones represent region B, while the central and the southern areas are summarized as region C. The distribution of blood sample numbers over the course of the years 2021 and 2022 is shown for each region in Figure 1. It becomes clear that there is a peak in sample collection during the summer months. This increase is mainly attributable to a rise in samples from juvenile birds during the summer, while the sample numbers from adult birds display no clear trend throughout the year. It is important to note, that one cooperation partner (Figure 1, No. 5) submitted blood samples in 2021 but not in 2022.

**Figure 1:**
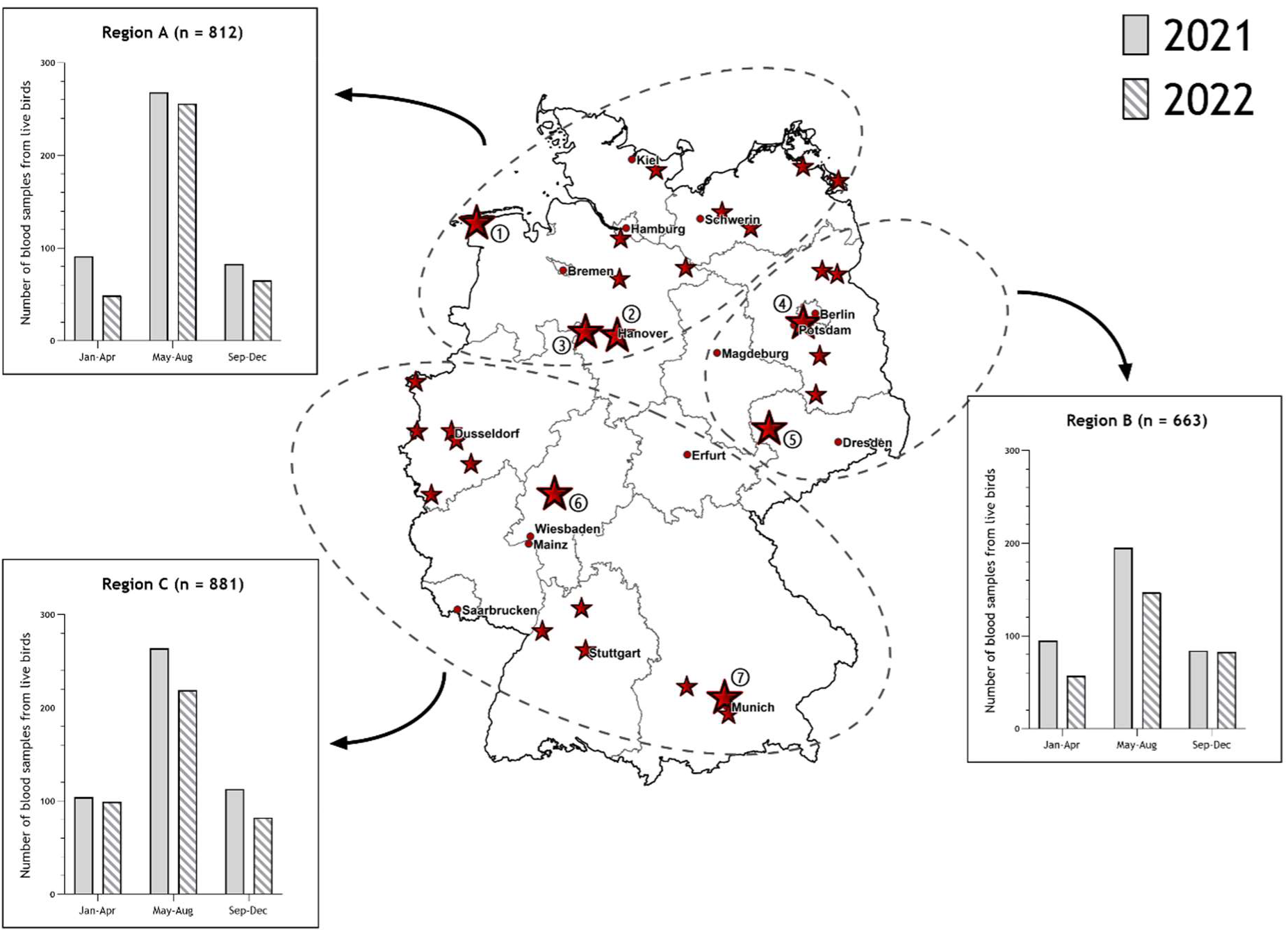
Locations of cooperation partners in the WBA-Zoo are depicted by big and small red stars and in some cases also by additional circled numbers (see below). Distribution of sample collection throughout 2021 and 2022 for each region is shown in separate graphs. The total number (n) of samples per region from both years combined is given in each graph title. Two samples from region B in 2021 are not shown in the corresponding bars as there are no exact dates of sampling available, but they are included in the aforementioned total number. Numbered sample collectors: 1. Seehundstation Nationalpark-Haus Norden-Norddeich, 2. Department of Small Mammal, Reptile and Avian Diseases, University of Veterinary Medicine Hanover, Foundation, 3. Wildlife Rescue and Conservation Centre Sachsenhagen, 4. Department of Veterinary Medicine, Small Animal Clinic, Freie Universität Berlin, 5. Clinic for Birds and Reptiles, Faculty of Veterinary Medicine, Leipzig University, 6. Clinic for Birds, Reptiles, Amphibians and Fish, Justus Liebig University Giessen, 7. Clinic for Birds, Small Mammals, Reptiles and Ornamental Fish, Centre for Clinical Veterinary Medicine, Ludwig Maximilians University Munich

#### 2.1.2 Organ samples (dead bird monitoring)

The second monitoring panel comprises mostly organ samples from 1791 deceased wild and captive birds from 2021 and 1318 individuals from 2022. The carcasses were submitted from locations distributed all over Germany (Supplementary Figure SF1). Then, organ samples were collected at bird clinics, zoos, wild parks and academic institutes, tested at the regional state veterinary diagnostic laboratories and subsequently submitted to the NRL. Many of these specimens were originally submitted to the regional laboratories for surveillance of avian influenza resulting in a larger share of waterfowl and captive birds from zoos or falconries, e.g. great grey owls, than in the live bird panel (Supplementary Table S4). Some of the birds sampled by cooperation partner No. 4 (Figure 1) while alive were submitted to the regional laboratory upon death. Therefore, results from blood and organ sample analysis are available for these birds (mainly Northern goshawks, *Accipiter gentilis*) and can be viewed in Supplementary Table S5.

The regional state veterinary diagnostic laboratories and the NRL as well as a cooperation partner from the Bernhard Nocht Institute for Tropical Medicine (BNITM, Hamburg) document their findings concerning presence of WNV and USUV in a shared West Nile Fever (WNF) database established by epidemiologists at the FLI (Friedrich-Loeffler-Institut).

### 2.2 Molecular Investigations

Viral RNA was extracted from cruor or organ material using the RNeasy Mini Kit (Qiagen, Hilden, Germany) according to the manufactureŕs instructions. The extracted RNA was routinely submitted to two reverse transcription quantitative real-time polymerase chain reaction (RT-qPCR) protocols for specific detection of USUV (Jöst et al. 2011) and WNV (Eiden et al. 2010). One protocol detects the non-structural protein 1 (NS 1) gene of USUV, the other one uses WNV-specific 5’ non-translated region (NTR) primers and probe. In accordance with the guidelines for the NRL, quantification cycle values (Ct) <37 were regarded as positive, from 37 to 40 as inconclusive and >40 as negative. In order to confirm the detection of USUV or WNV genomic viral RNA, blood samples yielding a positive or inconclusive result were additionally screened by a RT-qPCR for the NS5 gene of USUV (Cavrini et al. 2011) or a WNV RT-qPCR targeting the genomic NS2A region (Eiden et al. 2010), respectively. Positive controls with 10^3^ and 10^4^ copies of USUV or WNV genome per reaction were included in each assay.

### 2.3 Whole genome sequencing and Phylogenetic Analyses

A selection of USUV and WNV RNA positive cruor and organ samples was submitted to whole genome sequencing (WGS). These samples were taken from various locations across Germany in the case of USUV and from a variety of known circulation hotspots in the case of WNV (Supplementary Tables S6 and S7). In few cases, sequences could not be recovered due to low sample quality and/or low viral loads (Ct-values >30).

WGS of USUV and WNV was performed via amplicon sequencing on a MinION device (Oxford Nanopore-Technology (ONT), Oxford Science Park, UK) as a combined approach of two previously reported protocols by Holicki et al. (Holicki et al. 2022) and Santos et al. (Santos et al. 2023). Briefly, the viral nucleic acid from WNV or USUV positive samples was reverse transcribed into cDNA by means of the SuperScript IV First-Strand Synthesis System (Cat. No. 18091050; Invitrogen by Thermo Fisher Scientific, Darmstadt, Germany) using random hexamers as unspecific primers (Quick et al. 2017). Subsequently, double stranded DNA amplicons were generated by conventional PCR using AccuPrime Taq DNA Polymerase High Fidelity (Cat. No. 12346-086/-094; Invitrogen) as enzyme. For each virus two separate oligonucleotide primer sets (Oude Munnink et al. 2019; Grubaugh et al. 2019) enabled specific amplification of cDNA generated from viral nucleic acid, thereby creating overlapping amplicons in two separate reactions. This was followed by a purification step using AMPure XP SPRI reagent (Cat. No. A63881, Beckman Coulter, Inc., Indianapolis, US), which is the means of purifying intermediate nucleic acid products throughout the entire protocol. The library preparation commenced with pre-treatment of amplicon strand ends for barcode ligation via NEBNext Ultra II End Repair/dA-Tailing Module (Cat. No. E7546, New England Biolabs (NEB), Ipswich, US) followed by barcode ligation (NEBNext Ultra II Ligation Module, Cat. No. E7595, NEB and Native Barcoding Expansion Kit 1-12/13-24, Cat. No. EXP-NBD104/-114, ONT), motor protein ligation (NEBNext Quick Ligation Module, Cat. No. E6056, NEB) and the addition of loading beads (Ligation Sequencing Kit, Cat. No. SQK-LSK109, ONT). Spot-ON flow cells (R.9.4.1, ONT) were prepared for usage on a MK1c sequencing instrument (ONT) via Flow Cell Priming Kit (Cat. No. EXP-FLP002, ONT).

After the sequencing process, Fast5 raw data reads were base-called with high accuracy, demultiplexed and trimmed using the Mk1C sequencer (Guppy v6.5.7, ONT). Additional demultiplexing and adaptor removal were conducted using Porechop on the NanoGalaxy platform (Koning et al. 2020). The quality of sequencing data was assessed with NanoComp (Coster and Rademakers 2023). Consensus sequence generation was performed using k-mer alignment (KMA) (Clausen et al. 2018) and Minimap2 (Li 2018).

The alignment of sequences utilized the Clustal W algorithm, implemented in MEGA v.11 software (Tamura et al. 2021). The optimal model for nucleotide substitutions was determined using jModeltest v.2 (Darriba et al. 2012). Maximum likelihood trees were then reconstructed using PAUP* v.4 (Swofford 2003), employing the subtree-pruning-regrafting branch-swapping algorithm to search for the heuristic tree. The reliability of tree topologies was assessed through bootstrap testing (1000 replicates) and the finalized trees were reconstructed using FigTree v.1.4.3 (Rambaut 2016). For WNV lineage classification, we followed the workflow previously described by Santos et al. (Santos et al. 2023). In brief, the complete coding sequences were utilized with the APC algorithm and AHC included in the R package ‘apcluster’ v1.4.10 (Bodenhofer et al. 2011), implemented in R Studio (v2022.02.2-485).

### 2.4 Serological Investigations

The majority of the obtained sera was screened for presence of anti-flavivirus IgG antibodies (Ab) using a commercial blocking ELISA (Ingezim West Nile Compac, Ingenasa, Madrid, Spain) according to the manufactureŕs instructions. This assay enables species-independent detection of Ab against domain III of the envelope protein of WNV. The results were differentiated according to inhibition percentage (IP; ≥40% was regarded as positive, <40% and >30% as doubtful and ≤30% as negative). Recently, it has been demonstrated that this commercial ELISA is sufficiently able to detect cross-reactively binding USUV-specific Ab as well as WNV-specific Ab (Ziegler et al. 2022).

Samples yielding positive or doubtful ELISA results were further differentiated by virus neutralization tests (VNT) as described by Seidowski et al. (Seidowski et al. 2010) with minor modifications. They were tested against WNV lineage 2 (WNV-2), Germany (GenBank accession number MH924836) and USUV lineage Europe 3, Germany (HE599647). Positive and negative control sera were included in the VNTs. The positive controls stem from experimentally infected or hyperimmunized animals, mostly birds, and possess a well characterized neutralizing Ab (nAb) titre (ND_50_).

To determine the ND_50_ of the sera, the reciprocal of the serum dilution that could inhibit the cytopathogenic effect of the applied virus in >50% of its replicates was determined using the Behrens-Kaerber method (Mayr, A., Bachmann, P.A., Bibrack, B.,Wittmann, G. 1977). A serum sample was considered positive for USUV-specific nAb if the ND_50_ in the USUV VNT was equal to or higher than 10 and if additionally, the ND_50_ in the corresponding WNV VNT was negative (ND_50_ < 10) or significantly lower (= fourfold lower). WNV VNT results were scored in the same manner. All those serum samples that had been non-reactive in the blocking ELISA and consequently were not tested by VNT were considered negative for specific Ab against either virus.

As the aforementioned ELISA cannot differentiate Ab against USUV and WNV, low sample volumes of less than 30 µl were not pre-screened by this method, but submitted directly to the VNTs. Some cytotoxic or haemolytic serum samples had to be excluded from the evaluation due to persisting inconclusive results. Similarly, in the cases of four individuals where a double sampling became evident after serological analysis had already been performed on both specimens, only the sample taken at an earlier timepoint was included in the calculation of detection rates. Whenever nAb titres for USUV and WNV did not deviate sufficiently from each other to allow clear differentiation, the corresponding serum was also excluded from the calculation of nAb detection rates and received the status “infection by an unspecified flavivirus” (Supplementary Table S8). A slightly different manner of interpretation of VNT results was adopted for samples from region B, where USUV and WNV are known to be co-circulating. Here, sera with less than a fourfold difference between USUV and WNV nAb titres and a high titre of at least 100 in one of the VNTs were not classified as “infected by an unspecified flavivirus” but rather interpreted as belonging to birds that had undergone serial infections by USUV and WNV. These cases were included into the calculation of detection rates for USUV as well as WNV nAb. The findings from sera of migratory birds were also included in the calculation of these detection rates.

### 2.5 Statistical Analyses

R version 4.2.2 was used for the calculation of nAb detection rates and 95% confidence intervals (95% CI). Comparisons of nAb detection rates between age groups and months of the year were made using Fisher’s exact test. The same test was applied for comparison of virus RNA detection rates between years. The statistical analysis was conducted using SPSS software (IBM Corp. Released 2011, IBM SPSS Statistics for Windows, Version 20.0, IBM Corporation, Armonk, NY, USA). A p-value of less than 0.05 was considered to be statistically significant.

### 2.6 MAPS

ArcGIS ArcMap 10.8.2 (ESRI, Redlands, CA, USA) and open data from GeoBasis-DE/BKG 2023 were used in order to perform Geographic Information System (GIS) analysis of the sampling sites as well as the origin of WNV nAb positive birds and USUV RNA positive and negative birds.

### 2.7 Ethical Statement

Most of the blood samples (first monitoring panel) were either taken for diagnostic purposes at veterinary clinics, wild bird rescue or falconry centres, wildlife parks and zoos or immediately after euthanasia of birds with an infaust prognosis. Samples of sentinel ducks and nestlings were taken as part of animal experiments approved by the competent authority of the federal states of Mecklenburg-Western Pomerania (MV), Brandenburg (BB) and Berlin (BE), Germany (MV: 7221.3-2-006/19, 7221-3-2-003/21, V 244-26631/2019; BB: 2347-A-10-1-2019; BE: G 0144/21) on the basis of national and European legislation, namely directive 2010/63/EU on the protection of animals used for scientific purposes. Tissue samples (second monitoring panel) were only taken from deceased birds submitted for necropsy.

## 3 Results

### 3.1 RT-QPCR results

#### 3.1.1 Blood samples (live bird monitoring)

Out of 2356 blood samples collected in 2021 and 2022, cruor was available for 2333 of them. In detail, we analysed 1286 RNA isolates by RT-qPCR in 2021 and another 1047 specimens in 2022.

Presence of USUV RNA was confirmed for blood samples from the regions A and B in 2021 and from all three regions in 2022. The earliest and latest detection of USUV nucleic acid in the live bird panel in 2021 both occurred in blood samples collected near the North Sea coast (Figure 1, No. 1) in region A (beginning of August: European herring gull, *Larus argentatus*; mid-October: common wood pigeon, *Columba palumbus*). In the following season (2022), USUV genome was first detected in early June in a carrion crow (*Corvus corone*) sampled near Giessen (Figure 1, region C, No. 6) and finally in early October in a Eurasian blackbird from Munich (Figure 1, region C, No. 7). Eurasian blackbirds (n = 9) and common wood pigeons (n = 9) make up the majority of USUV RNA positive cases in this panel.

In both years, WNV-specific nucleic acid in blood samples was only detected in birds found in Berlin or Brandenburg (Figure 1, region B, No. 4). In the 2021 transmission season the earliest and the latest case of WNV infection were a Northern goshawk sampled in mid-July and a common wood pigeon sampled mid-September, respectively. In the following year (2022) the first evidence of WNV circulation was found in a European kestrel (*Falco tinnunculus*) in mid-July and the last one in a feral pigeon (*Columba livia f. domestica*) sampled at the beginning of November. Northern goshawks were the species most often infected by WNV in both years (n = 19). Many WNV RNA positive individuals of this predatory species showed unspecific as well as neurological clinical symptoms like apathy, ataxia, tremor and anisocoria. Several birds died during treatment or had to be euthanized and their carcasses were subsequently submitted to the responsible regional veterinary laboratory for further investigations. Therefore, laboratory results from molecular and serological investigations performed on blood as well as organ samples are available for 26 Northern goshawks and additionally two hooded crows (*Corvus cornix*, Supplementary Table S5).

According to the live bird panel there was an increase in USUV cases in 2022 (Table 2), which constitutes a significant difference (6/1286 in 2021 as opposed to 16/1047 in 2022, *p-value* < 0.05). Meanwhile, the numbers of WNV infections per year in the cohort from region B (13/288 and 28/267) did not differ from each other in a significant manner (Table 2, *p-value* ≥ 0.05).

**Table 2:**
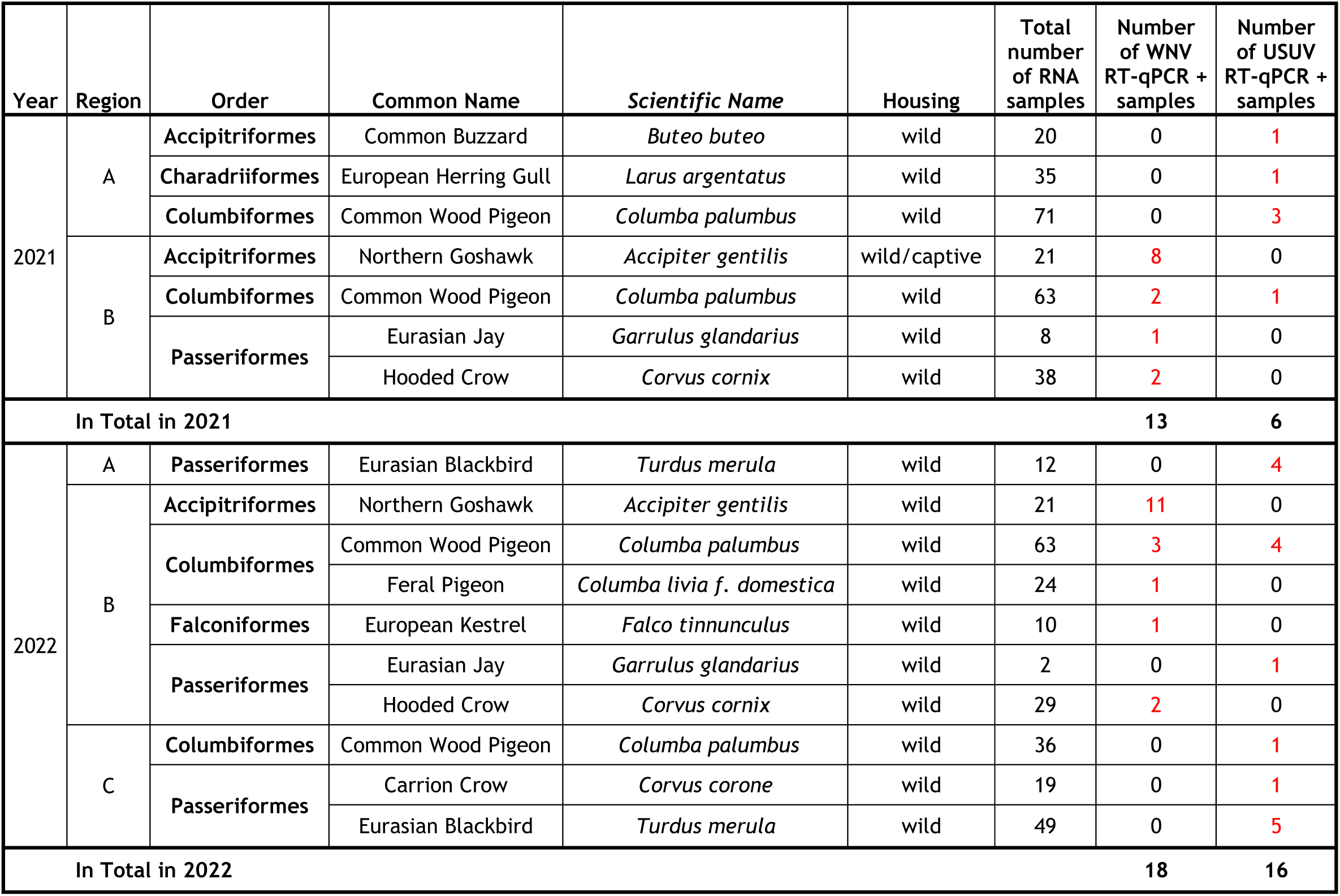
Positive (+) RT-qPCR results from blood samples (live bird panel) tested in 2021 (1286 RNA samples) and 2022 (1047 RNA samples). The relevant flavivirus is highlighted by red font colour. Background information on the affected species includes the total number of RNA samples from the same species tested by RT-qPCR in the respective region and year and the manner of housing.

#### 3.1.2 Organ samples (dead bird monitoring)

A higher rate of USUV infections in 2022 also became apparent in the dead bird panel (*p-value* < 0.05, Table 3 and Supplementary Figure SF1). Four autochthonous cases of WNV infection from outside region B (Figure 1) were submitted to the NRL in 2022 (Table 4), namely from Hamburg (one carrion crow and one blue tit, *Cyanistes caeruleus*) and Thuringia (one coconut lorikeet, *Trichoglossus haematodus,* and one kea, *Nestor notabilis*).

**Table 3:**
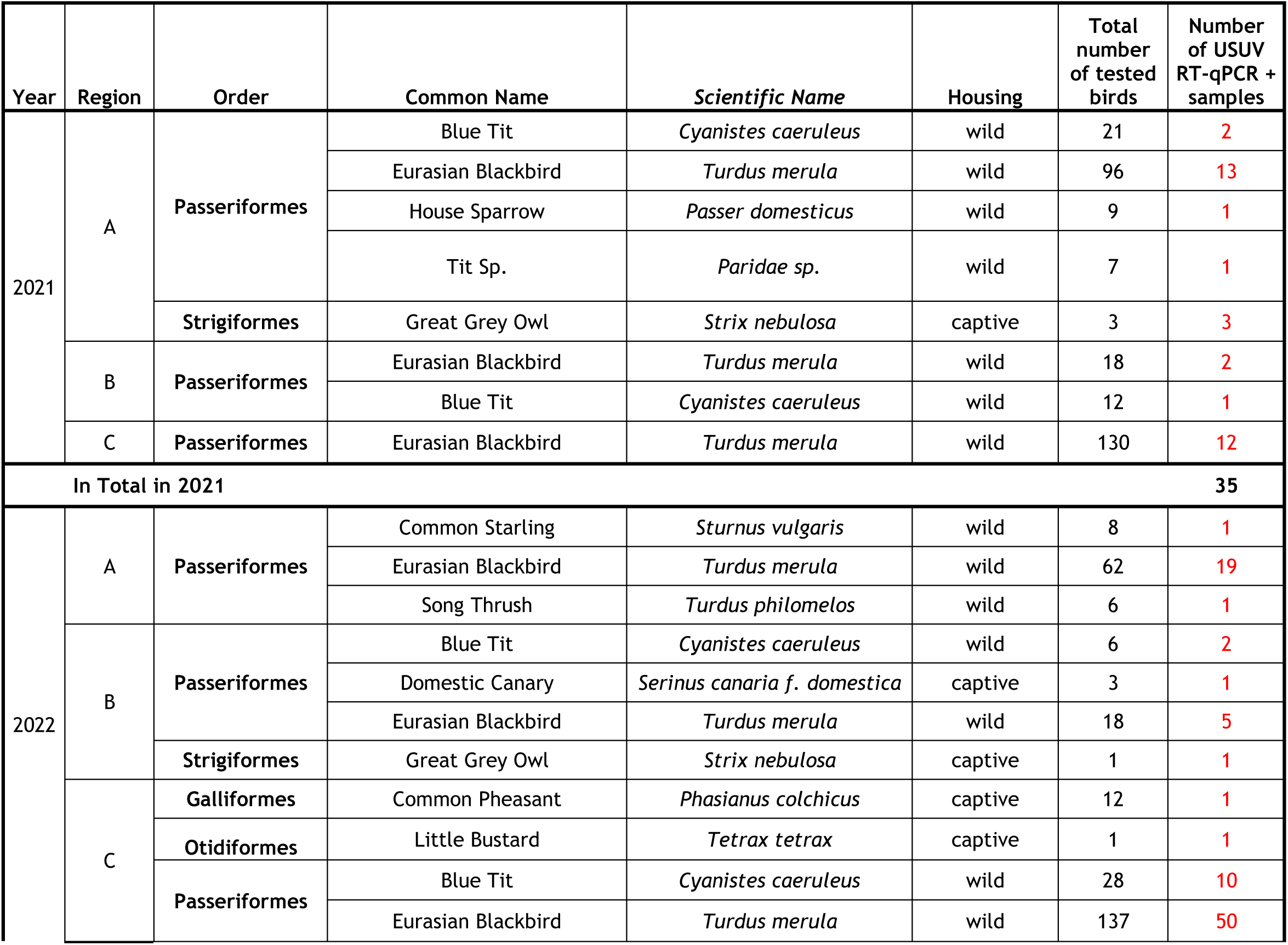
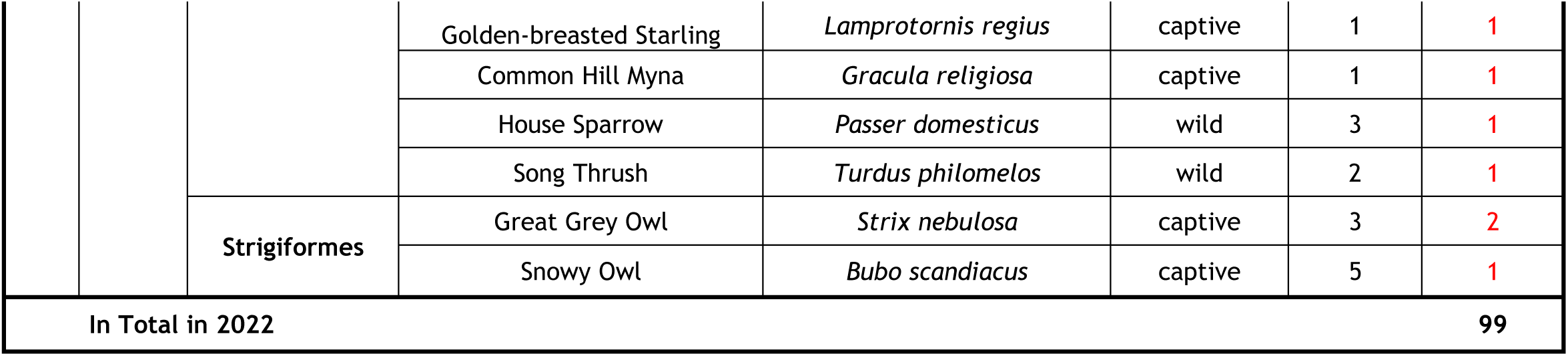
Positive (+) USUV RT-qPCR results from organ samples (dead bird panel) tested in 2021 (1791 RNA sample sets) and 2022 (1318 RNA sample sets). Background information on the affected species includes the manner of housing as well as the total number of birds from the same species tested by RT-qPCR in the respective region and year. Some USUV RNA positive cases from 2021 have also been published in a comparative study about the occurrence of avian influenza viruses and orthoflaviviruses in birds (Günther et al. 2023).

**Table 4:**
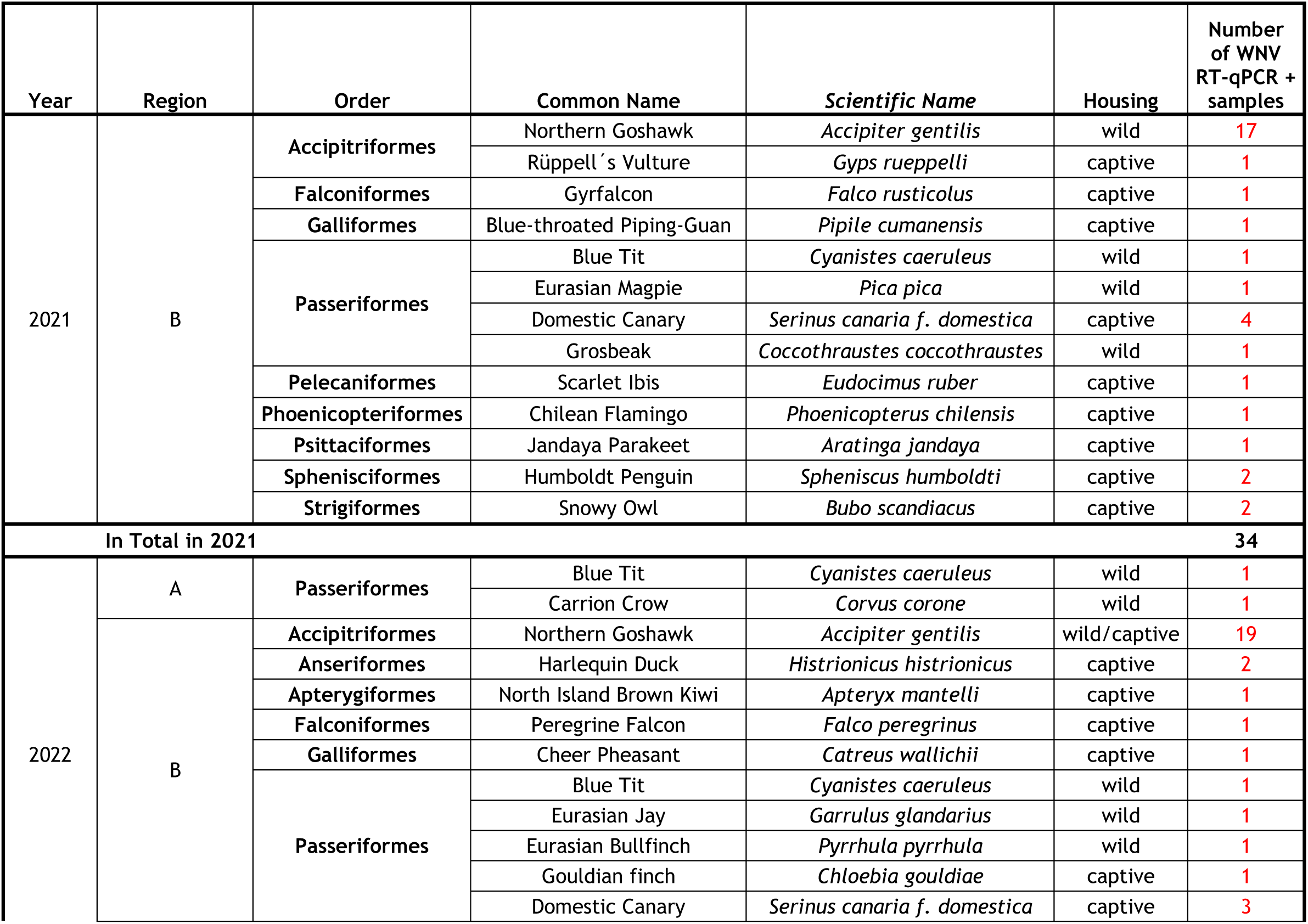
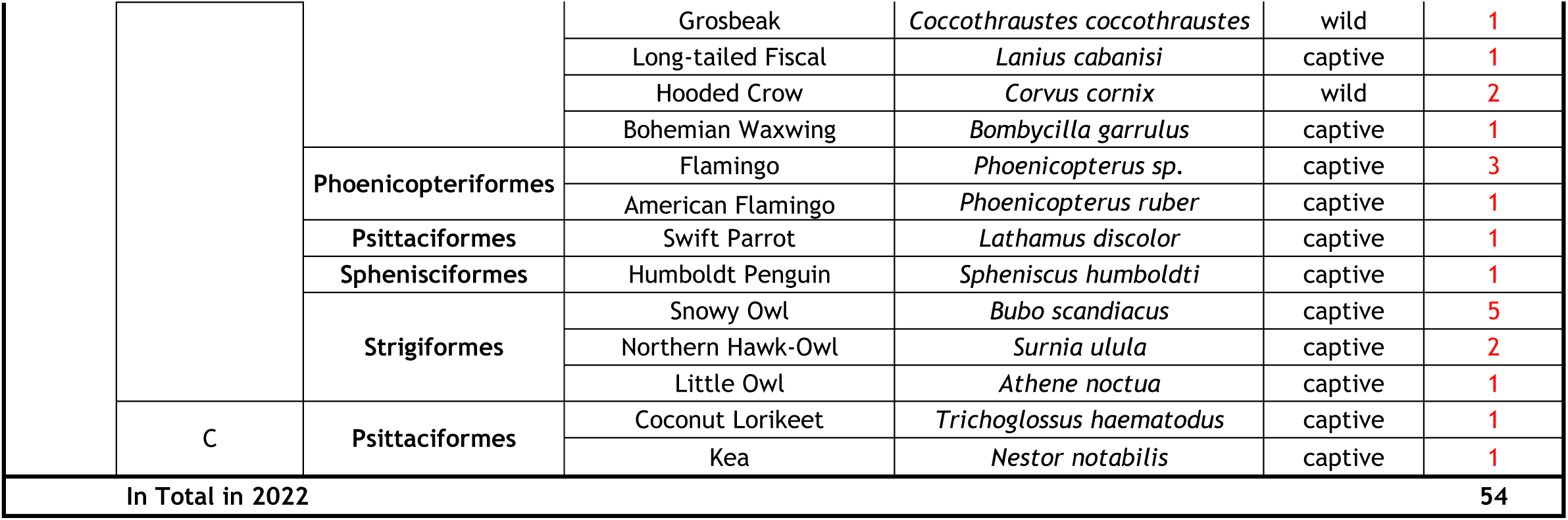
Positive WNV RT-qPCR results from organ samples (dead bird panel) confirmed in the NRL in 2021 and 2022. As WNV is a notifiable disease in Germany, positive cases are additionally registered in the German animal notification system (TSN database) and total numbers of tested birds may differ marginally from those in the FLI WNF database provided in Table 3. Background information on the affected species includes the manner of housing as well as the region of origin. Some WNV RNA positive cases from 2021 have also been published in a comparative study about the occurrence of avian influenza viruses and orthoflaviviruses in birds (Günther et al. 2023).

### 3.2 Phylogeny

Sequencing data of USUV RNA positive samples from the year 2021 have been published recently to portray the molecular evolution of USUV in Germany over the years (Bergmann et al. 2023a). Consequently, the sequencing panel for USUV consists only of samples from 2022 including two blood samples from the live bird panel as well as 13 organ samples from the dead bird panel.

For WNV, samples from both years were available for sequencing. In total, ten blood samples (2021: six samples, 2022: four samples) and 54 organ samples (2021: 27 samples; 2022: 27 samples) were sequenced.

#### 3.2.1 Phylogenetic analysis of USUV RNA positive birds (2022)

Overall, 15 USUV sequence samples from 2022 were analysed for phylogenetics and lineage distributions (Supplementary Table S6). The analysis revealed that seven samples were attributed to Europe 2 lineage, while four samples were classified under Europe 3 and another four samples were identified as belonging to Africa 3 lineage (Figure 2).

**Figure 2:**
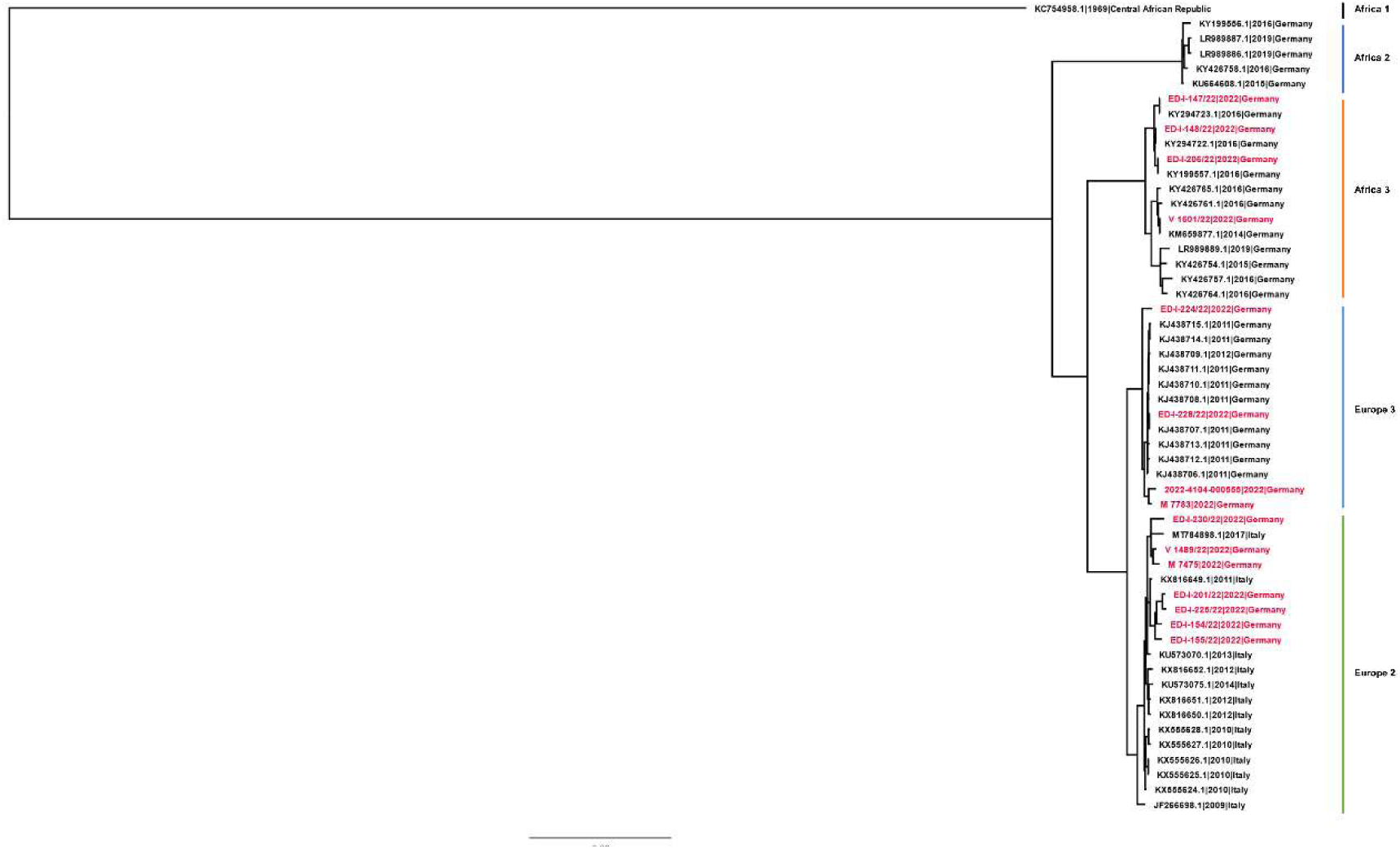
Phylogenetic analysis of USUV lineages. A maximum-likelihood tree was generated using complete genome sequences, with the scale bar indicating the mean number of nucleotide substitutions per site. The sequences are identified by codes that include the GenBank accession number, the sample collection year, and the country of origin. The sequences from Germany described in this paper are highlighted in red and GenBank accession numbers can be found in the supplementary file.

#### 3.2.2 Phylogenetic analysis of WNV RNA positive birds (2021 and 2022)

Phylogenetic analysis revealed that all WNV samples belonged to WNV-2. We employed clustering methods to identify the dominant subclade of WNV circulating in Germany during 2021 and 2022. Remarkably, 95% of the WNV sequences clustered within the 2.5.3.4.3c subcluster of WNV-2, including the WNV RNA positive carrion crow found dead in Hamburg in 2022 (Supplementary Table S7 and Supplementary Figure SF3).

Our analysis did not reveal any indication of new introductions of WNV strains into Germany during the study period. However, it is worth noting that three cases from 2022 were identified as belonging to the subcluster 2.5.3.4.3.a and to the cluster 2.5.3.2 (Figure 3 and Supplementary Figure SF3). No other European clusters or subclusters of WNV-2 were detected in this study (Supplementary Figure SF2).

**Figure 3:**
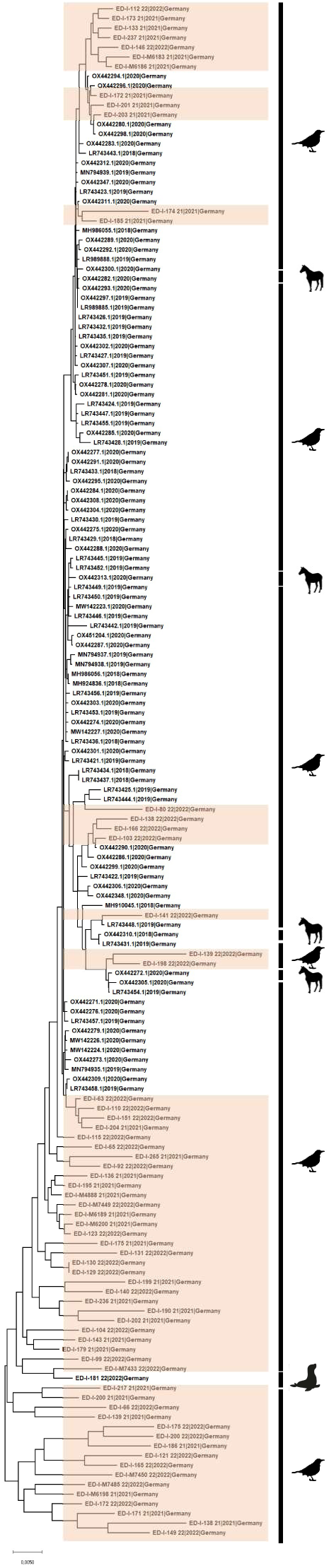
Phylogenetic analysis of lineage WNV 2. The maximumlikelihood tree was conducted based on complete genome sequences. Scale bar indicates mean number of nucleotide substitutions per site. Sequences are labeled by codes containing the GenBank accession number, year of sample collection and country of origin. Sequences acquired in this study are highlighted in orange and GenBank accession numbers can be found in the supplementary file. The pictograms on the right indicate the animal group of origin.

### 3.3 Serological results

From the 2356 wild birds tested in both years 2166 sera were available for diagnostic work-up (2021: 1229 sera, 2022: 937 sera). The percentage of samples reactive in the flavivirus-specific ELISA varied between regions (Figure 4). In region A 20.84% (134/643, 95% CI 17,76%-24,19%) of the tested samples showed a positive or doubtful result, in region B 46.38% (288/621, 95% CI 42,40%-50,39%) and in region C 14.78% (124/839, 95% CI 12,45%-17,36%).

**Figure 4:**
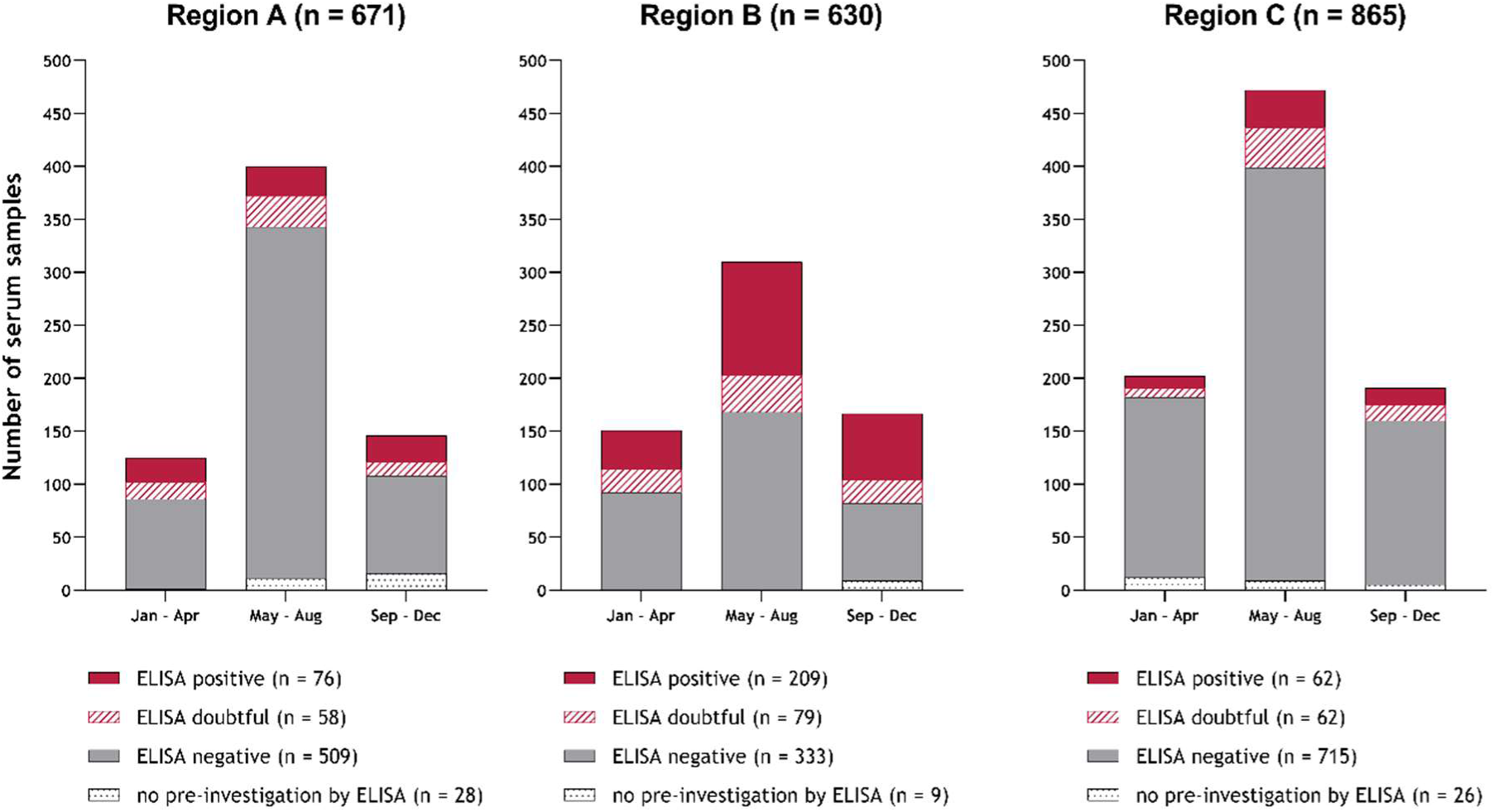
ELISA results and distribution of the investigated serum samples throughout the years 2021 and 2022 in regions A-C. The total number (n) of serum samples per region from both years is given above each graph, the number of serum samples with a specific ELISA outcome per region in the legend below, which elucidates the colour codes of the three possible ELISA outcomes (positive, doubtful, negative) and of the samples that were not pre-investigated by ELISA due to small sample volume. Two ELISA positive samples from region B in 2021 are not depicted in the corresponding graph as there are no exact dates of sampling available but they are included in the written numbers.

As this assay detects Ab against WNV as well as USUV a differentiation by separate WNV- and USUV-specific VNTs becomes necessary. The detection rates of WNV- and USUV-specific nAb for each of the three regions were calculated from the combined ELISA and VNT results. In both years, the USUV nAb detection rates were similar in all three regions (Table 5), while WNV nAb were clearly more abundant among birds from region B (Table 6). Serological indications of serial infections or potential co-infections were found once in a Eurasian blackbird in 2021 (USUV ND_50_/WNV ND_50_: 80/240) and four times in 2022 (two common buzzards, *Buteo buteo*, 320/100 and 160/100; one house sparrow, *Passer domesticus*, 120/320; and another Eurasian blackbird, 480/240). All of these individuals were sampled in region B (Figure 1).

**Table 5:** Detection rates of USUV nAb in blood sera (live bird panel) per region and year. The absolute numbers of seropositive and serologically tested birds are given in brackets (non-differentiated results excluded).

| Region/Year | Detection of USUV nAb | 95% CI |
| --- | --- | --- |
| A/2021 | 3.13% (12/383) | 1.63% - 5.41% |
| A/2022 | 3.24% (9/278) | 1.49% - 6.06% |
| B/2021 | 4.53% (16/353) | 2.61% - 7.26% |
| B/2022 | 7.23% (18/249) | 4.34% - 11.18% |
| C/2021 | 3.13% (15/479) | 1.76% - 5.11% |
| C/2022 | 4.70% (18/383) | 2.81% - 7.33% |

**Table 6:** Detection rates of WNV nAb in blood sera (live bird panel) per region and year. The absolute numbers of seropositive and serologically tested birds are given in brackets (non-differentiated results excluded).

| Region/Year | Detection of WNV nAb | 95% CI |
| --- | --- | --- |
| A/2021 | 1.04% (4/383) | 0.29% - 2.65% |
| A/2022 | 1.08% (3/278) | 0.22% - 3.12% |
| B/2021 | 17.85% (63/353) | 14.00% - 22.25% |
| B/2022 | 23.29% (58/249) | 18.19% - 29.05% |
| C/2021 | 1.25% (6/479) | 0.46% - 2.71% |
| C/2022 | 0.26% (1/383) | 0.01% - 1.45% |

### 3.4 Regional distribution of USUV antibody positive birds

In 2021, 43 of 1229 birds tested positive for nAb against USUV, in 2022 the number was 45 out of 937 individuals (Details in Supplementary Tables S9 and S10). The most commonly affected bird species in both years were Eurasian blackbirds (n = 21 out of 151 tested individuals from both years combined), common wood pigeons (n = 32 out of 311) and common buzzards (n = 13 out of 162).

The rate of juvenile birds sampled by our cooperation partners in relation to the size of their cohorts was very variable (Supplementary Table S11). This heterogeneity makes comparison of nAb detection rates between sites difficult. Therefore, only samples from adult birds were used to explore the situations at sampling site level (Figure 5 and Supplementary Table S12). An exception was sample collector No. 5 (Figure 5) in 2021 as no information on bird age was available. Therefore, the detection rate given for this location in 2021 includes all age groups and cannot be compared with the other sites.

**Figure 5:**
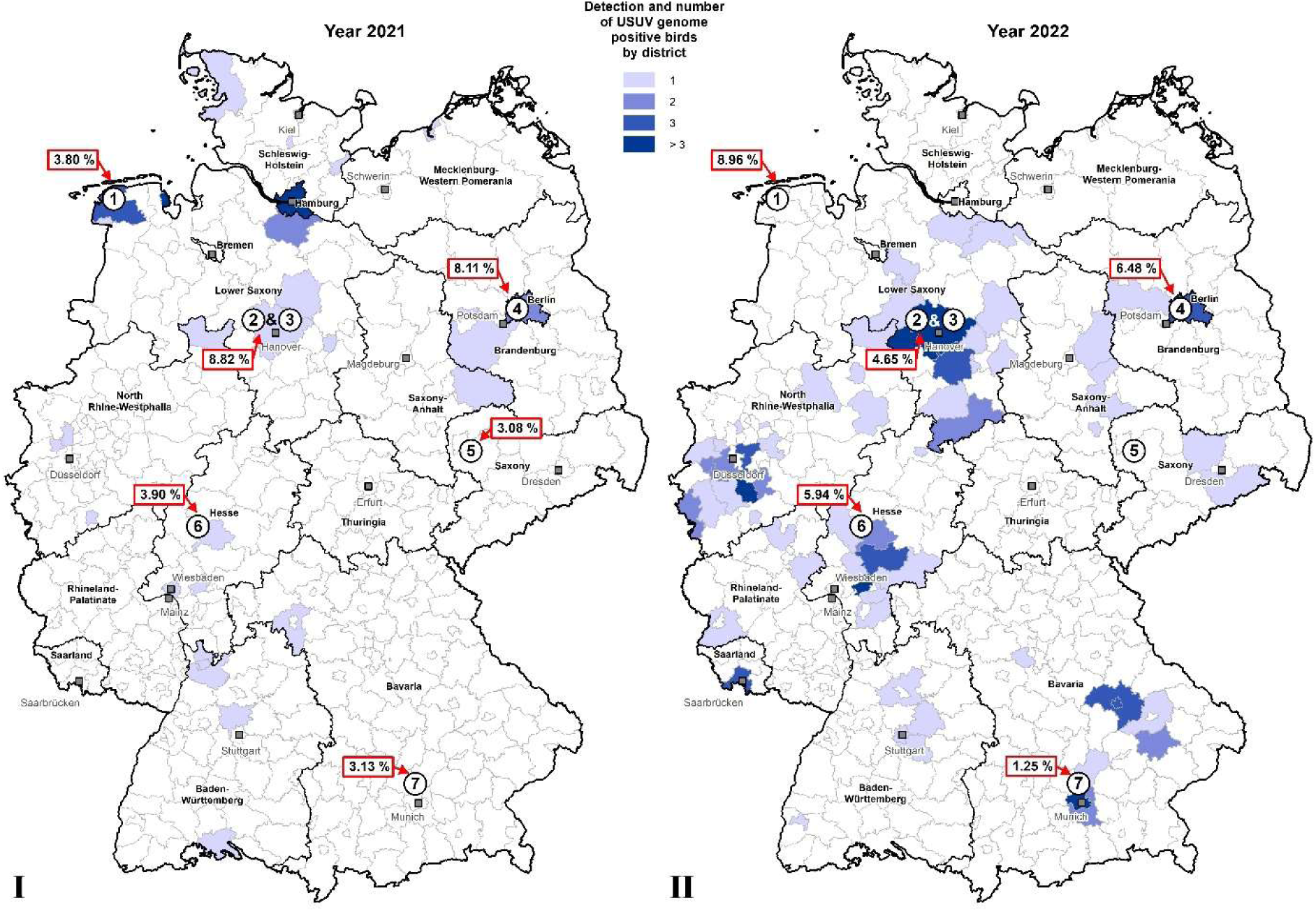
Detection rate of USUV nAb at selected sampling sites in Germany for the years 2021 (I) and 2022 (II). The circled numbers belong to the cooperation partners listed for Figure 1. Due to geographical overlaps regarding bird origin, the cohorts of sample collectors No. 2 and 3 were combined for the calculation of detection rates. Positive results from the live and dead bird panel in USUV-specific RT-qPCR for the respective year and at district level are indicated by different shades of blue in the background.

### 3.5 Regional distribution of wnv antibody positive birds

The cases of birds with nAb against WNV clustered in the German east with only sporadic occurrences outside of region B (Figure 6). Overall the sera from 73 out of 1229 birds tested in 2021 and from 62 out of 937 individuals in 2022 showed a positive WNV VNT result (Supplementary Tables S9 and S10). In region B, Northern goshawks (n = 33 out of 41 tested individuals from both years combined) and common wood pigeons (n = 28 out of 127) were the species with the highest total number of WNV nAb positive results. In total, 121 birds from region B displayed WNV-specific nAb. It is important to note, that in 2021 and 2022 some cases of birds with nAb against WNV occurred outside the enzootic hotspots in region B (Figure 1). More specifically, there were 14 cases of WNV nAb detected in region A (seven) and C (seven). Interestingly, one of these WNV nAb positive birds in 2021 was a great spotted woodpecker (*Dendrocopos major*) from near Munich in region C (Figure 1). Members of this species display strictly resident behaviour in southern Germany. The same is true for a seropositive carrion crow from Hanover in region A (Figure 1) in the year 2022. Other seropositive birds from regions A and C belong to species for whom at least short migratory behaviour cannot be completely excluded, but who predominantly live as resident birds or partial migrants (e.g. common buzzards or European kestrels). This means that the population in its entirety or at least parts of it stay in Germany year-round.

**Figure 6:**
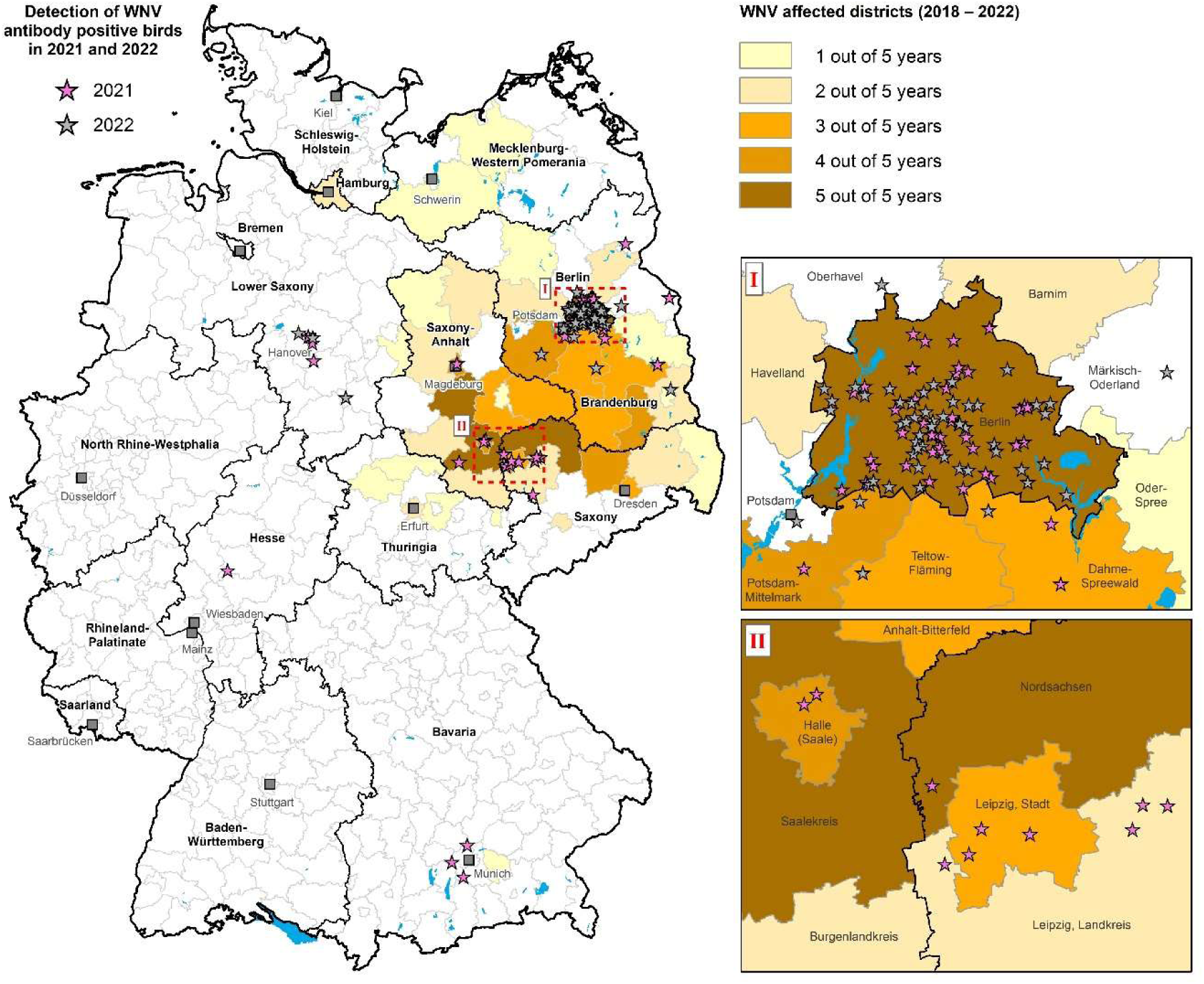
Origin of WNV nAb positive birds (resident and partially migratory) in 2021 and 2022 marked by pink and grey stars, respectively. The different shades of orange at district level indicate WNV-affected districts in Germany between 2018 and 2022 meaning that cases of WNV RNA positive birds or WNV RNA and/or IgM-Ab positive horses were registered. The hotspot areas around Berlin (I) and Leipzig (II) are displayed in more detail on the right.

### 3.6 Influence of age on the presence of nAb against USUV and WNV

In 2021 not one juvenile individual was found to possess specific Ab against USUV. In 2022, only one non-adult common wood pigeon found near Hanover and six young birds sampled in Berlin (Figure 5, No. 4) showed positive USUV VNT results. The totals of seropositive birds where the age is unknown amount to seven and five individuals from region B in 2021 and 2022, respectively. This means that the majority of samples with nAb against USUV came from adult birds (2021: 36 birds, 2022: 33 birds).

The outcome reveals a marked divergence in USUV nAb detection rates between juvenile and adult birds when looking at the total numbers per year (*p-value* < 0.05).

Comparable to the occurrence of USUV nAb, no juvenile bird from region A and only one such individual from region C in 2022 tested positive for nAb against WNV. In contrast, the WNV nAb detection rates of juvenile and adult birds sampled in the known WNV hotspot Berlin (Figure 5, No.4) show no significant difference (*p-value* ≥ 0.05, Table 7).

**Table 7:** WNV nAb detection rates per age group and year in the cohort sampled by cooperation partner No. 4 (Figure 5). The absolute numbers of seropositive and tested birds are given in brackets (non-differentiated results excluded).

| Age group | Year 2021 | 95% CI (2021) | Year 2022 | 95% CI (2022) |
| --- | --- | --- | --- | --- |
| Juvenile | 16.95% (20/118) | 10.67% - 24.96% | 23.36% (25/107) | 15.73% - 32.53% |
| Adult | 17.12% (19/111) | 10.63% - 25.43% | 24.07% (26/108) | 16.37% - 33.25% |
| Age unknown | 19.30% (11/57) | 10.05% - 31.91% | 21.88% (7/32) | 9.28% - 39.97% |

A close look at the occurrence of WNV nAb positive results over the course of 2021 and 2022 at this location shows that the peak in serum sample collection from juveniles appears in July. However, the maximum of WNV seropositive juveniles is reached in August, approximately two to six weeks after the first WNV infections detected in cruor by RT-qPCR in this study (Figure 7, graphs a) and b)). The increase in WNV nAb detection in juvenile birds from July to August, followed by a decrease from August to September, is remarkable for both years (*p-value* < 0.05). Meanwhile, there is no notable variation concerning the detection of WNV nAb throughout the year among adult birds in region B (Figure 7, graphs c) and d)).

**Figure 7:**
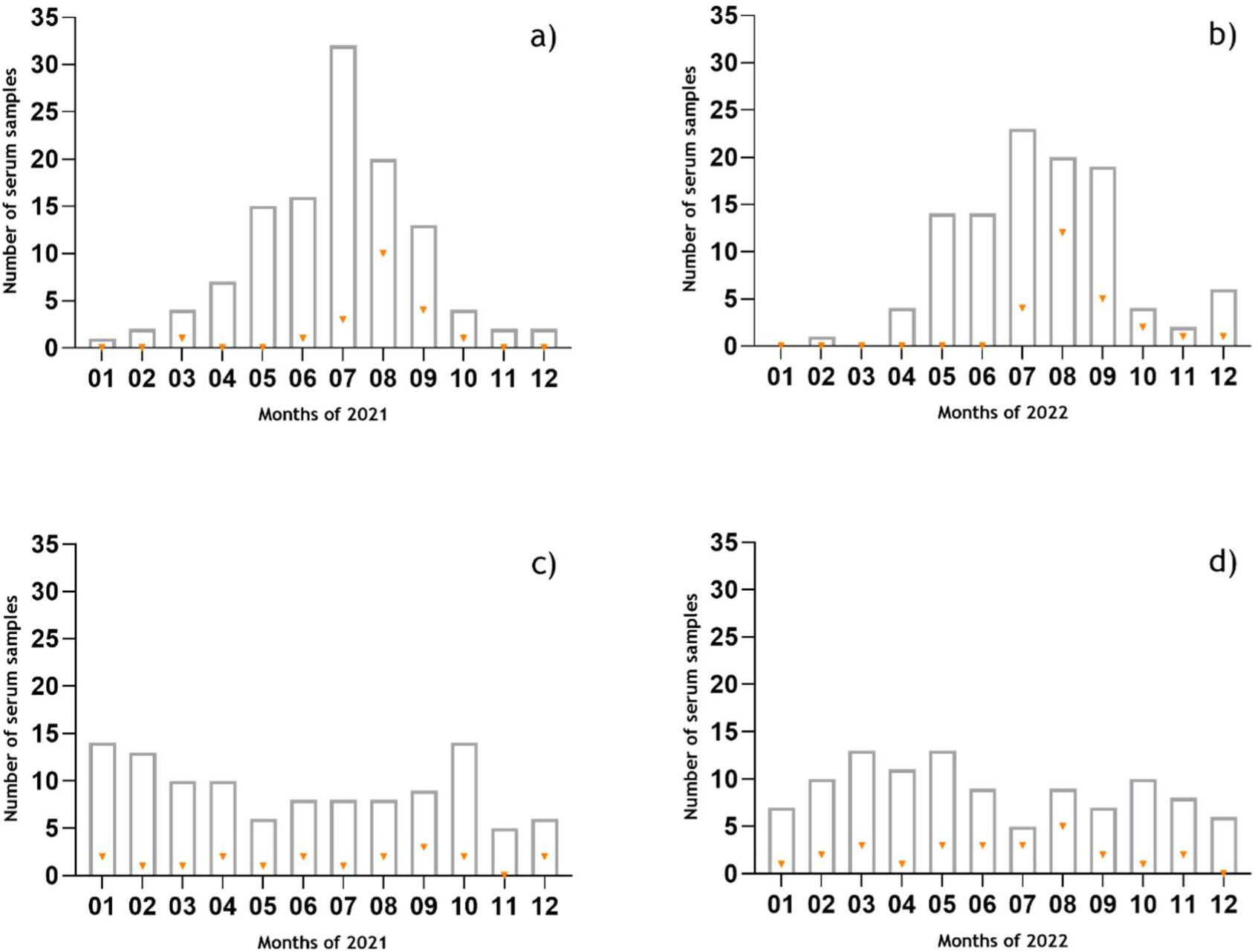
Relation between the number of serum samples tested per month and the number of WNV nAb positive outcomes (Sample collector No. 4, region B, Figure 1). The total number of tested samples for each month is indicated by the bars, while the orange triangles mark the number of WNV seropositive cases. Graphs a) and b) represent the juvenile population in 2021 and 2022, respectively, while the results for the adult birds are displayed in the graphs c) and d). The first WNV RNA positive cases of the seasons at location No. 4 were detected on 15.07.2021 and 14.07.2022, respectively.

Sample collector No. 4 (Figure 5) additionally identified some of the juvenile birds as “pullus”. From both years combined, sera from five of these extremely young chicks showed a positive result in a WNV VNT, one bird was USUV nAb positive and six sera were reactive but could not be differentiated by VNT (Supplementary Table S13). All of these seropositive birds belonged to pigeon sp., most were common wood pigeons. There were two instances of acute WNV infection in pigeon pulli in 2021 (Supplementary Table S13, birds No. 2 and 4) revealed by RT-qPCR matching the positive result of the corresponding WNV VNT. In another case (Supplementary Table S13, No.6), the bird tested positive for USUV by RT-qPCR, while the nAb found in its serum were directed against WNV. The majority of all nAb titres in this age group was relatively low (between 10 and 30 ND_50_) with only three exceptions (Supplementary Table S13, No. 4, 7 and 9).

## 4 Discussion

While USUV-specific RNA was found in dead and wild birds throughout the country, the detection of WNV genomes was almost exclusively limited to region B (eastern and central-eastern Germany, Figure 1). This is consistent with WBA-Zoo findings from preceding years (Michel et al. 2019; Ziegler et al. 2022).

For USUV, periodical surges in infection rates have become apparent since its first detection in a German bird in 2011. This pattern reappeared once more in 2022 as a notable increase in USUV RNA positive cases is evidenced by our results in the live bird as well as dead bird panel for this year after comparatively low infection rates in 2020 and 2021 (Ziegler et al. 2015; Michel et al. 2018; Michel et al. 2019; Ziegler et al. 2022). Nonetheless, the number of USUV RNA positive cases did not reach the level of the year 2018, when Germany in its entirety was affected for the first time with more than 1000 detected avian infections (Ziegler et al. 2019a). As expected, genomic RNA of USUV in blood samples from live birds was mainly detected in summer and early fall of 2021 and 2022.

The recurring surges in USUV genome positive cases may be partly attributable to a reversed ebb and flow in population immunity. When looking at the samples from adult birds, the yearly USUV nAb detection rates at the sampling locations show different trends. Overall, a herd immunity against USUV as described by Meister et al. (Meister et al. 2008) for a region in Austria in the year 2005 has not been reached at any of the monitored locations.

The USUV whole genomes sequences from the year 2021 (dead bird panel, n=5) were published previously by Bergmann et al. (Bergmann et al. 2023a). Four of them belonged to USUV lineage Africa 3 and one to Europe 3. While the consistent identification of lineages Europe 2, Europe 3 and Africa 3 in our USUV sequence panel of 2022 underscores the enduring presence and circulation of diverse USUV lineages within Germany, Europe 2 now represents almost half (7/15) of the whole genome sequences generated from the live and dead bird panel in 2022. In 2019 and 2020, the detection rate of this lineage amounted merely to 6/37 and 4/25 specimens, respectively. Of particular interest is the observed emergence of Europe 2 lineage cases in the federal states of Lower Saxony and Bavaria, where this lineage had not been detected before. The potential transmission of Europe 2 lineage strains from eastern and central-eastern Germany (Figure 1, region B), where they were previously prevalent, to other regions highlights the interconnectedness of USUV transmission pathways within Germany (Michel et al. 2019; Ziegler et al. 2022; Bergmann et al. 2023a). Further investigations into the drivers behind the (re-)emergence of specific USUV lineages are essential for a comprehensive understanding of viral circulation dynamics and the development of targeted surveillance and control strategies.

Similar to the development of USUV infections, the live bird panel also displayed a rise in WNV genome positive cases in 2022. The difference turned out to be not statistically significant, but the trend coincides with the dynamics in other European countries like Italy. In this Mediterranean country, the number of human WNV cases during the 2022 transmission season were the highest ever reported and the number of animal outbreaks was only surpassed by the ones recorded in 2018 (European Food Safety Authority and European Centre for Disease Prevention and Control 2023).

As already mentioned, avian WNV infections in contrast to USUV remain restricted to certain regions of Germany. A high WNV nAb detection rate of more than 20% in 2022 is only reached in region B (Figure 1, Table 6). When looking at the geographical distribution of WNV nAb positives in our study (Figure 6), however, it becomes evident that there are some cases of WNV immunity from other areas of the country. These could be signs of an enzootic or silent circulation of WNV between reservoir hosts (birds) and vectors (mosquitoes) as suggested by Zehender et al. (Zehender et al. 2017) after phylogenetic analysis of a large quantity of WNV-2 isolates from all over Europe. Zehender and colleagues showed that the introduction of the virus in e.g. Hungary, Austria, Greece and Italy took place several years before animal and human outbreaks were detected. The most blatant indicators of such non-apparent infection events in our study are the WNV nAb positive great spotted woodpecker sampled near Munich (2021) and the carrion crow sampled near Hanover (2022) as these species are considered to be almost exclusively resident. Therefore, these birds have to have undergone WNV infection locally, most probably following a mosquito bite. No acute WNV infections in birds or mammals have been reported from the area around Hanover so far. Furthermore, the last confirmed WNV cases from near Munich date back to 2018, when two great grey owls from a wild park (Poing) died due to WNV (Ziegler et al. 2019b; Ziegler et al. 2020). Emergence of West Nile disease in humans and animals in these areas may be imminent as was the case in Hamburg in the summer of 2022, where in addition to the carrion crow and the blue tit from our dead bird panel one horse was diagnosed with acute WNV infection (Friedrich-Loeffler-Institut and Schweizerisches Bundesamt für Lebensmittelsicherheit und Veterinärwesen 2022). For this area, first serological evidence of WNV introduction was found in a juvenile zoo bird from a wild park south of Hamburg (Bergmann et al. 2023b). Bearing in mind the serological signs of silent virus circulation at several locations, it was also no great surprise, when in the summer of 2023 a juvenile snowy owl (*Bubo scandiacus*) was diagnosed with acute WNV infection in a previously unaffected German federal state in the south west (Rhineland-Palatinate) (Friedrich-Loeffler-Institut and Schweizerisches Bundesamt für Lebensmittelsicherheit und Veterinärwesen 2023).

Nonetheless, the overall slow geographical spread of WNV through Germany since its introduction in 2018 mirrors the situation in other European countries. It took neuroinvasive WNV-2 approximately four years to bridge the distance between eastern Hungary, its place of initial detection (2004), and Austria (2008). Meanwhile, a neuroinvasive WNV-1 strain managed to cross the entire North American continent within the same time span after its emergence on the eastern coast in 1999 (Nash et al. 2001; Glaser 2004; Bakonyi et al. 2006; Bakonyi et al. 2013). One reason for the difference in expansion velocity could be cross-protective immunity to other flaviviruses within the European bird population. Co-circulation of WNV and USUV for example has been shown for many European countries in addition to Germany, e.g. Hungary, Austria, the Netherlands, Italy and Greece (Simonin 2024). Interestingly, Reemtsma et al. (Reemtsma et al. 2023) observed that a preceding USUV infection protected experimentally WNV infected geese from severe disease. Viremia and shedding of WNV were decidedly reduced in comparison to experimental WNV monoinfection (Reemtsma et al. 2022). A partial protection against severe WNV disease could also be demonstrated for Eurasian magpies (*Pica pica*), which were previously infected with USUV (Escribano-Romero et al. 2021). Therefore, the observed USUV seropositivity in the wild bird population in Germany, although the percentage is variable and on a low level, could be one of several explanations for slower geographical spread of WNV within the country.

In both years, the earliest WNV RNA positive sample from a wild bird from location No. 4 (Figure 1) had been taken more than one week before the first human case in Berlin was diagnosed (09.08.2021 and 02.08.2022, (European Centre for Disease Prevention and Control 2022, 2023)). This qualifies wild bird monitoring as an early-warning system for human health once more.

The Northern goshawk appears to be of special importance for WNV circulation at sampling location No. 4 (Figure 1). It is the species with the highest total number of RT-qPCR positive cases despite a decidedly lower total number of samples in comparison to e.g. common wood pigeons. The same is true for the presence of WNV nAb in members of this species. Ever since the emergence of WNV-2 in Hungary in 2004, several studies from European countries have noticed the high WNV infection rate among Northern goshawks often accompanied by severe neurological symptoms marking these birds as indicators of WNV circulation (Erdélyi et al. 2007; Hubálek et al. 2019; Vilibic-Cavlek et al. 2019). Several explanatory approaches have been suggested, e.g. the host feeding preference of mosquitoes for raptors in general in comparison to smaller birds (Victoriano Llopis et al. 2016) or the additional risk of alimentary infection existing for avian predators (Garmendia et al. 2000; Komar et al. 2003; Nemeth et al. 2006). In this context, special attention should also be paid to the feral pigeon (live bird panel) diagnosed with WNV infection via RT-qPCR as late in the year as November 2022. As pigeons and doves make up a considerable portion of the diet of Northern goshawks (Engler et al. 2021), this might lead to alimentary infection of these raptors even during the winter months. This was for example suggested for one WNV-2 infected Northern goshawk with neurological symptoms from the province of Perugia, Italy, that died in January 2022 (Mencattelli et al. 2022). The transmission of WNV from prey to predator may be an additional though rare overwintering strategy for the virus during the break in mosquito activity in temperate zones complementing the mechanisms of virus persistence in hibernating mosquitoes (Vidaña et al. 2020; Kampen et al. 2021; Mencattelli et al. 2022). Heightened awareness for the relevance of Northern goshawks in WNV epidemiology could have led to slightly biased sampling in our study setup resulting in a slight overrepresentation of the species in our sample panels. For truly unbiased assessment of the WNV susceptibility of Northern goshawks in general and via the oral route an animal trial would be necessary, which can be no easy feat to accomplish for a wild bird species this large. For now, this leaves careful interpretation of monitoring data as the best source of knowledge on infection dynamics in this species.

The consistent identification of WNV-2 across all samples in this study indicates the stable predominance of this viral lineage within Germany (Santos et al. 2023). The clustering analysis revealed a predominant subcluster, 2.5.3.4.3c, comprising 95% of the sequenced WNV strains, suggesting a high degree of genetic relatedness and indicating the dominance of a specific viral sublineage. This clustering pattern and the absence of indicators for new introductions suggest ongoing local transmission and circulation of genetically similar strains within the German population of WNV vectors and hosts for the years 2021 and 2022. Meanwhile, the identification of cases belonging to other clusters and subclusters highlights the enduring presence of genetic diversity within the circulating strains, emphasizing the need for continued surveillance to monitor potential shifts in viral sublineages and the introduction of novel strains (Santos et al. 2023).

The differentiating view on juvenile and adult birds provided in this study is new to the WBA-Zoo. This has led to several interesting novel insights into orthoflavivirus epidemiology. Nationwide, we found only seven cases of USUV nAb in juveniles. Meanwhile, the detection rates of WNV nAb in juveniles and adults from sample collector No. 4 (Figure 1) are almost identical and amount to about 20% (Table 6). As per definition juveniles in our study have experienced less than two complete WNV/USUV transmission seasons, this would mean that about one fifth of the sampled juvenile birds at location No. 4 (Figure 1) get infected by WNV within this time span. While potential persistence of Ab from previous transmission seasons complicates the use of serological data from adult birds as a surveillance tool for virus circulation (Gibbs et al. 2005), results gained from juvenile serum samples may indeed provide helpful insights. This has already been shown for domestic pigeons (*Columba livia f. domestica*) in Greece (Chaintoutis et al. 2014). As portrayed in Figure 7, our data clearly show a rise and subsequent fall in WNV seropositive cases among juvenile birds during the summer months of both years with a peak reached in August, two to six weeks after the first RT-qPCR positive cruor sample from the live bird panel. Allowing time for the onset of Ab production, this seems to be a direct response to heightened virus circulation. Further studies would be necessary to evaluate the use of this dynamic as an early-warning indicator for virus circulation. In this context, one has to bear in mind the possibility of passive transfer of immunity rather than active immunization as has already been shown for WNV-1 and USUV (Gibbs et al. 2005; Hahn et al. 2006; Meister et al. 2008). The detection of WNV nAb in pulli from Berlin (Figure 1, region B, No. 4) indicates the same phenomenon for WNV-2 infections.

To conclude, signs of inapparent enzootic circulation of WNV in Germany and recent outbreak scenarios of WNV in other European countries with high numbers of human infections and even fatalities stress the need for continued disease surveillance. In this context, the WBA-Zoo makes good use of available resources at minimum cost and remains an important source of knowledge on the infection dynamics of WNV and USUV in the wild bird population of Germany.

## Supplementary Materials

Figure SF1: Origin of tested dead birds from the dead bird panel (database) for 2021 (A) and 2022 (B), Figure SF2: Bayesian tree representing the time-scaled phylogeny of Germany’s WNV Subclade 2.5.3 complete coding sequences, Figure SF3: Geographical origin of WNV- sequences generated for the years 2021 and 2022, Table S1: Detailed register of the tested blood samples by bird orders and species from region A (northern and central-western part of Germany) from 2021 and 2022, Table S2: Detailed register of the tested blood samples by bird orders and species from region B (eastern and central-eastern part of Germany) from 2021 and 2022, Table S3: Detailed register of the tested blood samples by bird orders and species from region C (central and southern part of Germany) from 2021 and 2022, Table S4: Numbers of organ samples from the dead bird panel itemized according to year of sampling and taxonomic order of the species of origin, Table S5: Results from birds, of whom blood as well as organ samples were tested (Figure 1, sample collector No. 4), Table S6: Detailed information on the origin of phylogenetically analysed USUV from wild and captive birds in the live and dead bird monitoring in 2022, Table S7: Detailed information on the origin of phylogenetically analysed WNV from wild and captive birds in the live and dead bird monitoring in 2021 and 2022, Table S8: Samples for which no differentiation between WNV and USUV by neutralization assays from wild bird blood samples was possible in regions A–C from the year 2021 and 2022, Table S9: WNV- and USUV-positive neutralization assay results from wild bird blood samples from regions A–C in 2021, Table S10: WNV- and USUV-positive neutralization assay results from wild bird blood samples from regions A–C in 2022, Table S11: Number of serum samples per age group and year submitted by a selection of cooperation partners based on cohort size, Table S12: Details on USUV nAb detection rates in adult birds per sample collector and Table S13: Seroreactive pigeon pulli sampled at the FU Berlin with corresponding nAb titres and RT-qPCR results

## Acknowledgements

We thank the veterinary authorities and laboratories of the German federal states for providing us with samples, and we are grateful for their continued support as well as for the conscientious registration of samples in the FLI WNF database. We are also thankful to all contributors from the nationwide wild bird surveillance network in diverse bird clinics/practices, zoos and wildlife rescue and rehabilitation centres as well as falconry centres for taking and sending samples for the present study, especially Frank Seifert, Silvia Urbaniak and Marina Grebe (Raptor Rehabilitation Centre Rhineland), Claudia and Max Hartl (Raptor Centre and Wildlife Park Hellenthal), Stefan Rebscher (Raptor Research Centre Burg Guttenberg, Neckarmühlbach), Rachel Hein and Jonas Heck (Clinic for Birds and Reptiles, Faculty of Veterinary Medicine, Leipzig University), Marc Engler (NABU Wild Bird Rehabilitation Centre, Berlin), and the members of the Association to Support Veterinary Medicine in Free-ranging Birds (Verein zur Förderung der Vogelmedizin e.V.), Giessen. We highly value the aid provided by Patrick Wysocki in creating the maps in this manuscript and by Mathias Merboth in maintaining the FLI WNF database. A very special thank you is owed to Cora M. Holicki (FLI) for her efforts in teaching the sequencing methods used in this study. Last but not least we are grateful for the excellent technical assistance provided by Katja Wittig and Cornelia Steffen (FLI), Sabine Schiller (IZW), Katrin Erfurt and Jana Schömburg (Institute of Virology at the Faculty of Veterinary Medicine in Leipzig).

## Authorś contributions

Conceptualization, M.H.G., U.Z.

Methodology, F.S., B.S., F.B., M.K., K.H., A.S., C.F., R.L., U.Z.

Sample collection and preparation, F.S., F.B., D.F., R.R., K.M., A.G., A.Gl., R.S., V.G.,

M.R., L.F., O.K., M.Ri., K.S., V.S., K.H., R.L., U.Z.

Sample analysis, F.S., F.B., K.H., A.S., R.L., U.Z.

Data curation, F.S., B.S., F.B., D.F., R.R., K.M., A.G., A.Gl., M.K., R.S., V.G., M.R., L.F., O.K., M.Ri., K.S., V.S., K.H., A.S., C.S., R.L., U.Z.

Validation, F.S., B.S., F.B., K.H., R.L., U.Z.

Statistical testing, F.S., B.S., U.Z.

Resources, F.S., B.S., F.B., D.F., R.R., K.M., A.G., A.Gl., M.K., R.S., V.G., M.R., L.F., O.K.,

M.Ri., K.S., V.S., K.H., A.S., C.F., C.S.-L., C.S., R.L., J.S.-C., F.Br., M.L., R.K., T.W.V., M.H.G., U.Z.

Writing – original draft, F.S., B.S., U.Z.

Writing – review and editing, F.S., B.S., F.B., D.F., R.R., K.M., A.G., A.Gl., M.K., R.S., V.G., M.R., L.F., O.K., M.Ri., K.S., V.S., K.H., A.S., C.F., C.S.-L., C.S., R.L., J.S.-C., F.Br., M.L., R.K., T.W.V., M.H.G., U.Z.

Visualization, F.S., B.S., U.Z.

Supervision, B.S., M.K., M.H.G., U.Z.

