## Supplementary material for "Circulation of West Nile Virus and Usutu Virus in Birds in Germany, 2021 and 2022": Figures

**Supplementary Figure 1:** Origin of tested dead birds from the dead bird panel (database) for 2021 (A) and 2022 (B). The coloured circles indicate the origin of birds that tested positive (orange) or negative (green) for presence of USUV-specific RNA. The icons are placed randomly within the corresponding postal code area or if the former is not available the district of origin.

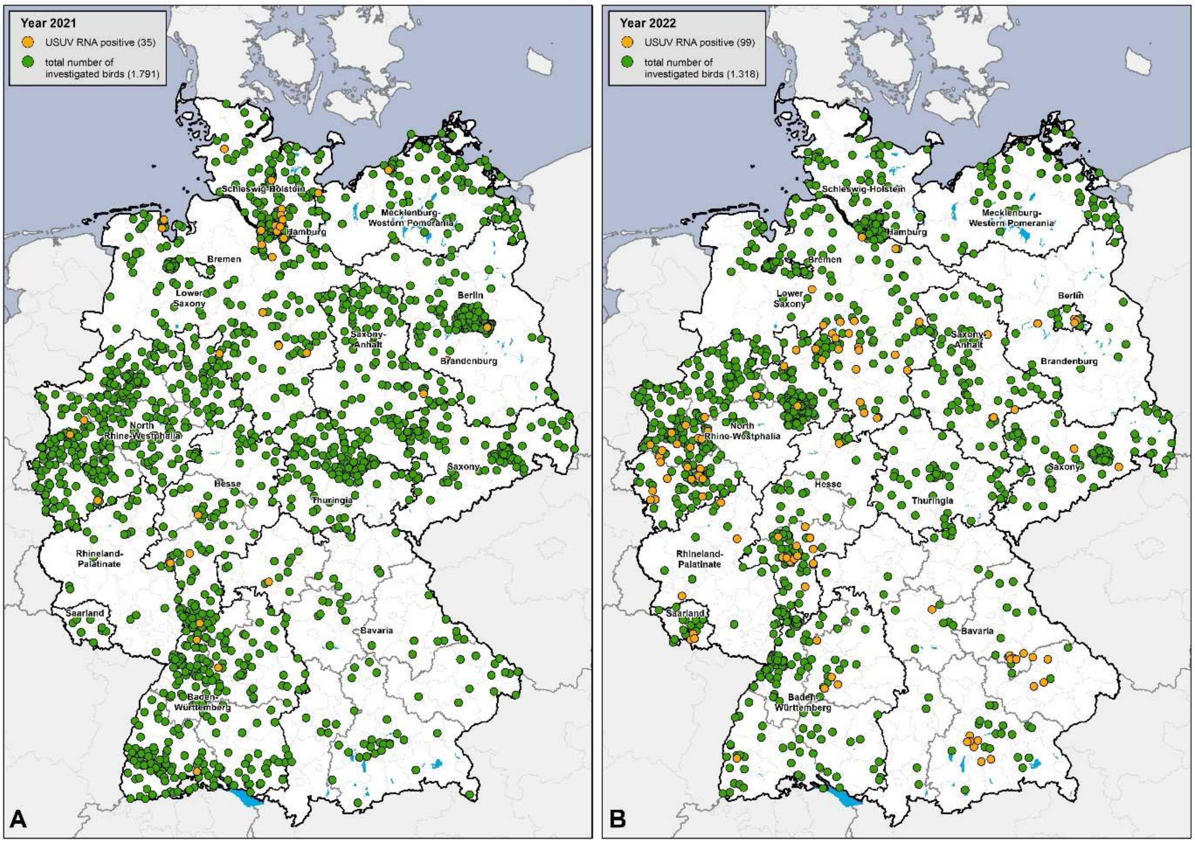

**Supplementary Figure 2:** Bayesian tree representing the time-scaled phylogeny of Germany's WNV Subclade 2.5.3 complete coding sequences. WNV sequences acquired in this study are highlighted in orange.

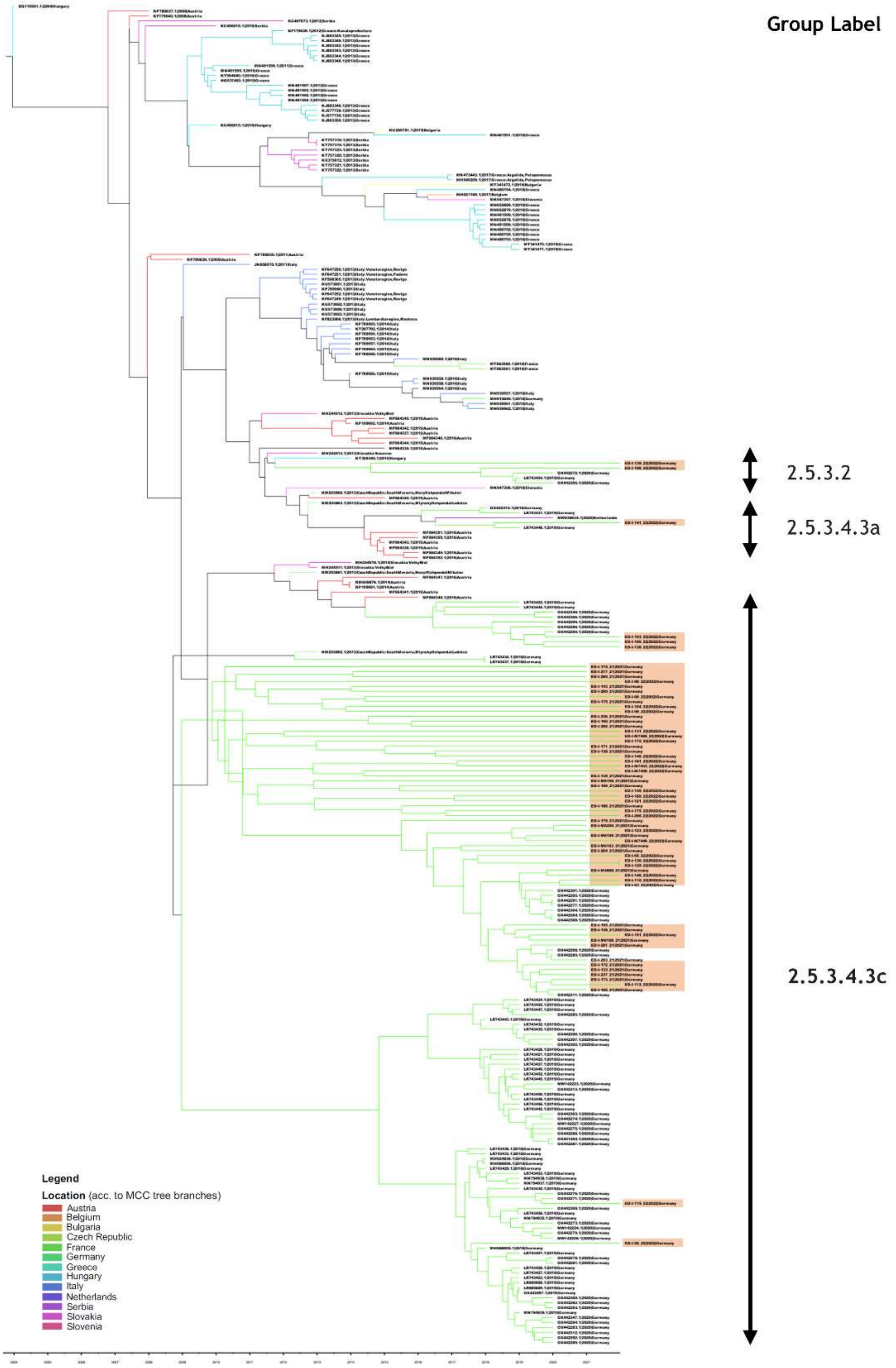

**Supplementary Figure 3:** Geographical origin of WNV-sequences generated for the years 2021 and 2022. The coloured icons mark the geographical origin of WNV sequences recovered from avian tissue in this study in 2021 (left) and 2022 (right). The different colours stand for the corresponding cluster or subcluster of WNV lineage 2. The sample material originated from wild and captive birds as indicated by icon shapes (see legend). The grey icons show the origin of WNV RNA-positive birds from the dead bird panel that were not submitted to sequencing.

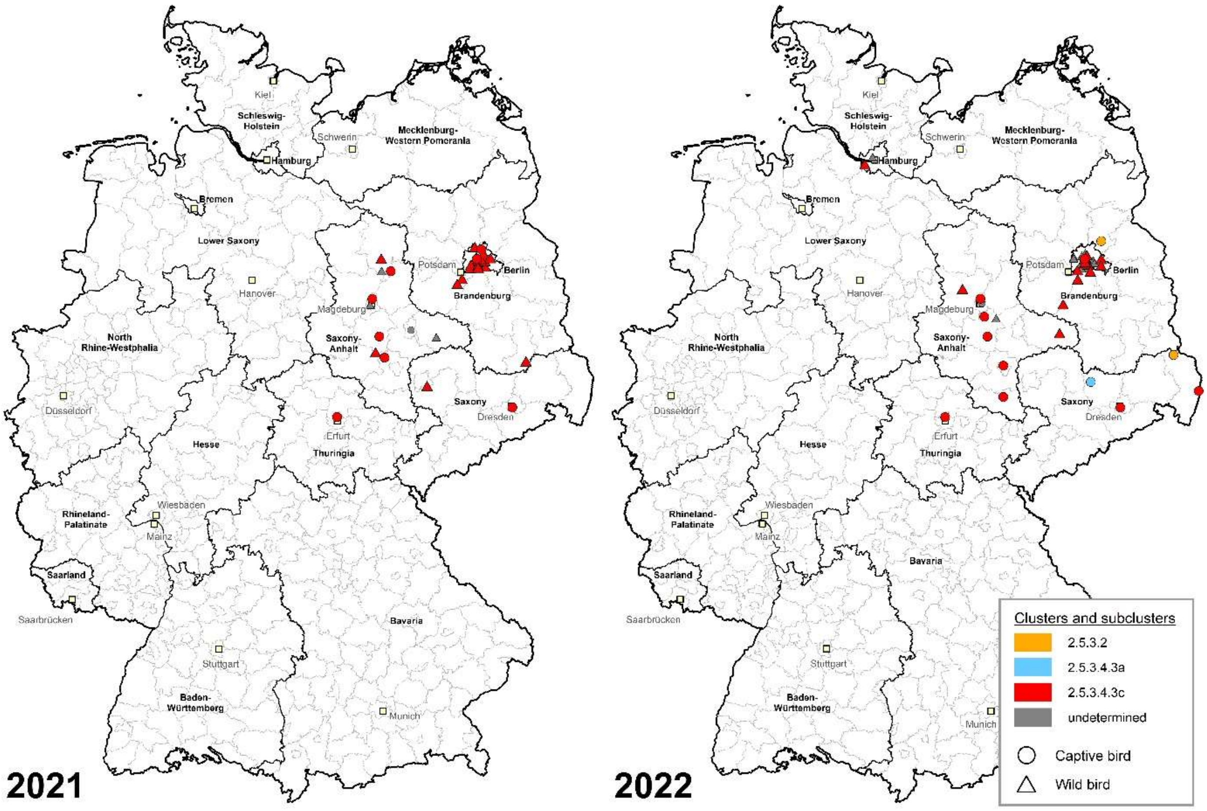
